# Identification of hub molecules of FUS-ALS by Bayesian gene regulatory network analysis of iPSC model: iBRN

**DOI:** 10.1101/2021.01.07.425798

**Authors:** Masahiro Nogami, Mitsuru Ishikawa, Atsushi Doi, Osamu Sano, Takefumi Sone, Tetsuya Akiyama, Masashi Aoki, Atsushi Nakanishi, Kazuhiro Ogi, Masato Yano, Hideyuki Okano

## Abstract

Fused in sarcoma/translated in liposarcoma (FUS) is a causative gene of amyotrophic lateral sclerosis (ALS). Mutated FUS causes accumulation of DNA damage and cytosolic stress granule (SG) formation, thereby motor neuron (MN) death. However, key molecular aetiology remains unclear. Here, we applied a novel platform technology, iBRN, “Non-biased” <u>B</u>ayesian gene regulatory <u>n</u>etwork analysis based on <u>i</u>nduced pluripotent stem cell (iPSC)-derived cell model, to elucidate the molecular aetiology using transcriptome of iPSC-derived MNs harboring FUS^H517D^. iBRN revealed “hub molecules”, which strongly influenced transcriptome network, such as miR-125b-5p-TIMELESS axis and PRKDC for the molecular aetiology. Next, we confirmed miR-125b-5p-TIMELESS axis in FUS^H517D^ MNs such that miR-125b-5p regulated several DNA repair-related genes including TIMELESS. In addition, we validated both introduction of miR-125b-5p and knocking down of TIMELESS caused DNA damage in the cell culture model. Furthermore, PRKDC was strongly associated with FUS mis-localization into SGs by DNA damage under impaired DNA-PK activity. Collectively, our iBRN strategy provides the first compelling evidence to elucidate molecular aetiology in neurodegenerative diseases.

**Highlights:**

- A new platform technology, “iBRN”, Bayesian gene regulatory network analysis based on iPSC cells
- iBRN identifies hub molecules to strongly influence the gene network in FUS-ALS
- PRKDC specifically acts as a guardian against FUS mis-localization during DNA damage stress
- miR-125b-5p-TIMELESS axis regulates DNA repair-related genes in FUS-ALS.

## INTRODUCTION

Amyotrophic lateral sclerosis (ALS) is a fatal neurodegenerative disorder characterized by selective death of upper (corticospinal) and lower somatic motor neurons (MNs) and about 10% of ALS patients have a familial history of the disease, categorized as familial ALS (fALS) (Brown and Al-Chalabi, 2017). The RNA-binding protein (RBP), Fused in sarcoma/translated in liposarcoma (FUS) was identified as an ALS causal gene. FUS protein contains a glycine-rich region, an RNA recognition motif, and a nuclear localization signal (NLS). Currently, more than 50 mutations of the FUS gene have been identified in fALS (Lattante et al., 2013, Vance et al., 2009). FUS protein is mainly localized in the nucleus (Anderson and Kedersha, 2009) but can shuttle between the nucleus and the cytoplasm (Dormann and Haass, 2011). However, the mutated FUS proteins in ALS patients are partially or totally excluded from the nucleus and form cytoplasmic inclusions in the neurons and glial cells of the brain and spinal cord (Dormann et al., 2010, Neumann et al., 2009, Suzuki et al., 2012). Interestingly, the ALS-linked mutations in FUS are mainly clustered at the C-terminal region of the protein, which contains the NLS. Indeed, increasing levels of cytoplasmic FUS caused by C-terminal mutations in FUS gene are associated with disease severity, early onset and fast progression of ALS (Dormann et al., 2010, Bosco et al., 2010, DeJesus-Hernandez et al., 2010). On the other hand, FUS has been implicated in the DNA damage response (DDR) signaling, induced by DNA double-strand breaks (DSBs). In response to DSB-inducing agents, FUS is phosphorylated by DNA-dependent protein kinase (DNA-PK), which are activated by DSBs (Han et al., 2012, Deng et al., 2014, Monahan et al., 2017, Murray et al., 2017, Rhoads et al., 2018). FUS interacts with histone deacetylase 1 (HDAC1) which may indirectly modulate DSBs repair by homologous recombination (HR) and non-homologous end joining (NHEJ). Furthermore, loss of FUS diminished both HR and NHEJ efficiencies (Wang et al., 2013). In addition, impairment of poly(ADP-ribose) polymerase (PARP)-dependent DDR signaling due to mutations in the FUS NLS, which caused cytoplasmic FUS aggregation and DNA nick ligation defects has been linked to ALS-related neurodegeneration (Naumann et al., 2018, Wang et al., 2018). However, the molecular mechanism of disturbed DDR caused by mutated FUS has not been fully understood yet. To date, several studies using human induced pluripotent stem cells (iPSCs) harboring different FUS mutant proteins have shown the cytoplasmic distribution of FUS proteins in human iPSC-derived MN (Ichiyanagi et al., 2016, Lenzi et al., 2015, Liu et al., 2015, Naujock et al., 2016). In addition, the enhanced recruitment of FUS proteins into cytosolic stress granules (SGs) after induction of cellular stress was observed (Aulas et al., 2012, Bentmann et al., 2012, Vance et al., 2013). In our previous study, we generated iPSCs from two fALS patients carrying the FUS^H517D^ mutation and healthy volunteers, as well as isogenic iPSCs homozygous for the FUS^H517D^ mutation, using the TALEN genome editing system, and differentiated these iPSCs into MNs to investigate their multifaceted cellular phenotypes in vitro (Ichiyanagi et al., 2016). We observed several neurodegenerative phenotypes including mis-localization of FUS protein to cytosolic SGs under stress conditions, and cellular vulnerability. The results suggested that our iPSC-derived MNs could be a useful tool to analyze the pathogenesis of human MN disorders.

Recent advances of systems biology using high throughput platforms such as microarrays and RNA sequencing has promoted understanding of complex biological and molecular disease aetiology. Among them, Bayesian gene regulatory networks (BRN) analysis are especially powerful bioinformatics approach for “non-biased” inference of causal relationships between mRNAs in those RNA abundance data sets to holistically model interactions between molecules in cells and tissues (Friedman et al., 2000, Imoto et al., 2002, Affara et al., 2007, Hartemink et al., 2001). In theory, BRN can elicit information about the complicated cellular regulatory relationships and a small number of “hub genes”, which are connected to large numbers of downstream “peripheral genes” in the networks, as candidate master-regulators of transcription and other cellular processes.

Here, we applied iBRN, which is a BRN analysis combined with iPSC-derived cell models to uncover molecular aetiology of FUS-dependent fALS (FUS-ALS). While the traditional bioinformatics revealed that DDR-related genes were uniformly down-regulated in iPSC-MN harboring mutated FUS. iBRN also identified well known DDR regulators, miR-125b-5p, TIMELESS and PRKDC (DNA-PK catalytic subunit) as top hub molecules which can strongly influence the gene regulatory network. Indeed, miR-125b-5p was up-regulated in the disease cell model and caused down-regulation of DDR-related genes including TIMELESS, PARP1 and FUS. Importantly, introduction of miR-125b-5p and reduced Timeless expression also caused the accumulation of DNA damage. Furthermore, the in vitro cellular model analyses validated the introduction of DSBs under impaired DNA-PK activity accelerated the cytosolic FUS mis-localization. Collectively, iBRN would be the “next generation” strategy to elucidate key molecular aetiology using iPSC-derived cell models in neurodegenerative diseases.

## EXPERIMENTAL PROCEDURES

### iPSC-derived motor neuron differentiation and immunofluorescence microscopic analysis

Motor neuron differentiation of iPSCs was performed as previously described with slight modification (Imaizumi et al., 2015, Ichiyanagi et al., 2016, Fujimori et al., 2018). Briefly, iPSCs were exposed to a medium including 3 μM SB431542 (SB), 3 μM dorsomorphin, and 3 μM CHIR99021 (CHIR) for over 5 consecutive days. Next iPSCs were detached from feeder layers and then enzymatically dissociated into single cells. The dissociated cells were cultured in suspension at a density of 1 × 105 cells per ml in low attachment culture dishes in MPC induction medium (media hormone mix; MHM) supplemented with 2% B27 supplement, 20 ng/ml FGF, 10 ng/ml hLIF, 2 μM SB, 3 μM CHIR, 2 μM retinoic acid (RA), and 1 μM purmorphamine (PM) in a hypoxic and humidified atmosphere (4% O2, 5% CO2) for 7 days. Next, neurospheres were passaged by dissociation into single cells and then cultured in MHM supplemented with 2% B27 supplement, 2 ng/ml bFGF, 10 ng/ml hLIF, 2 μM SB, 2 μM RA, and 1 μM PM for 7 days under 4% O2 hypoxic conditions. 5 μM DAPT was added to the modified MPC medium after 4 days from sphere passage (DIV16). The medium was changed every 2-3 days for a total of 14 days to induce MPCs. To differentiate neuronal cells, dissociated MPCs were plated onto 6-well plate or 96-well plates with poly-l-ornithine (PO) and growth-factor-reduced Matrigel (50 × dilution, thin coated; Corning) and cultured in differentiation medium that consisted of MHM supplemented with 2% B27 supplement, 10 ng/ml rhBDNF, 10 ng/ml rhGDNF, 200 ng/ml ascorbic acid, 1 μM RA, and 2 μM DAPT for 14-28 days in a humidified atmosphere (20% O2, 5% CO2. Half of the medium was changed every 2 or 3 days. For the immunofluorescence microscopy of iPS cells-derived neuronal cells, the cells were cultured on matrigel-coated 96-well, Black/Clear, Tissue Culture Treated Plate, Flat Bottom with lid (FALCON; #353219). After fixing with 4 % paraformaldehyde for 30 min at room temperature. The cells were then treated with 1 % bovin serum and 1 % goat or 1 % donkey serum, and 0.1 % Triton X-100 for 60 min at room temperature and incubated with mouse monoclonal anti-TUBB3 (Sigma; #T8660) (1:1000), mouse monoclonal anti-HB9 (DSHB; clon#81.5C10-c) (1:125), chicken polyclonal anti-CHAT (AVES; #CAT) (1:500) in PBS with 0.05% Tween-20 overnight at 4°C. After 3-times wash with PBS (-) at room temperature, the cells were incubated with Alexa Fluor 488 Goat Anti-chicken IgY, Alexa Fluor 555 Goat Anti-mouse IgG1 and Alexa Fluor 647 Donkey Anti-mouse IgG (H+L) secondary antibodies and 5 μg/ml Hoechst 33258 in PBS with 0.05% Tween-20 for 60 min at room temperature. Next, cells were washed with PBS (-) at 3 times. The immunofluorescence signals were observed under a fluorescence microscope (Keyence; BZ-9000) equipped with ×20 (Nikon; PlanApo, NA=0.75) objective lens.

### Large-scale transcriptome analysis of iPSC-derived MNs with a GeneChip HTA2.0

Purification of total RNA from the differentiated MNs was performed with miRNeasy Mini Kit (QIAgen). Total RNA quality was analysed using an Agilent 2100 BioAnalyzer (Agilent Technologies). Large-scale transcriptome analysis of 60 total RNA samples was performed with a GeneChip HTA2.0 (Affymetrix) based on the manufacturer’s user manuals, including a GeneChip WT PLUS Reagent Kit User Manual, GeneChip Expression Wash, Stain and Scan User Manual for Cartridge Array, GeneChip Fluidics Station 450/250 User Guide, and GeneChip Command Console (AGCC) 4.0 User Manual to generate CEL files, and then Expression Console Software 1.4.1 User Manual to generate CHP files. The obtained CHP files were analysed with Transcriptome Analysis Console Software (Affymetrix). A PCA was performed with the CEL files using Affymetrix Expression Console software. For GO analyses, DEGs (FDR P-Value < 0.05) in the comparisons were applied to GO analysis with DAVID Bioinformatics Resources 6.8 (https://david.ncifcrf.gov/). The data discussed in this publication have been deposited in NCBI’s Gene Expression Omnibus (GEO) and are accessible through GEO Series accession number GSE118336.

### miRNA profiling with an nCounter miRNA analysis system

miRNA profiling analysis of total RNA was performed with an nCounter Analysis System and a Human miRNA Assay Kit Version 3.0 (NanoString Technologies) using the FOV max mode. Data analyses were performed with nSolver analysis software version 2.0 equipped with the nCounter Analysis System. Normalization of miRNA profiling data was performed with the top 100 miRNA expression levels.

### iBRN analysis

BRN analysis (Friedman et al., 2000, Imoto et al., 2002) for estimating a gene regulatory network from high-throughput gene expression data has been introduced from two decades ago and used in the field of bioinformatics (Margolin et al., 2006, Huynh-Thu et al., 2010). The structure of Bayesian network can be represented as a directed acyclic graph (DAG) resulted from the process of estimation with many parameters which are estimated from given observed data matrix and correspond to find optimal structure of the graph. SiGN-BN is gene network estimation software that implements Bayesian network modelling and parameter estimation for high performance computing (Tamada et al., 2011). SiGN-BN automatically and appropriately estimated hyper parameters, starting from random variables by using the bootstrap method, a widely applicable and extremely powerful statistical technique which allows estimation of the sampling distribution using random sampling methods (RJ, 1994, H, 2005). For gene list selection in Bayesian network modelling, we selected several sets of interesting genes that includes about one thousand genes because SiGN-BN has a limitation the number of genes it can estimate. The iBRN analysis can be applied to each gene list. In the estimated structures of the network, there are nodes and edges which indicate the genes and the connection line between nodes with regulatory direction, respectively. The genes which have many edges with parental regulatory direction are called as “hub” genes and the hub genes may have a potential to affect many genes (Affara et al., 2007). Thus, the genes in our networks were sorted and allocated by the number of parental edges they had. For this study, the gene networks were estimated from 60 microarrays that were the same dataset of DEGs and GO analyses by the SiGN-BN software on Super-Computer with 1000 bootstrapping. The tree (Sugiyama) layout and organic layout were used to draw the gene networks. Super-Computer resource was provided by the Human Genome Center (The University of Tokyo, Tokyo, Japan). SiGN-BN gene network estimation software is currently available for Human Genome Center Super-Computer, RIKEN Center for Computational Science K computer and/or its compatible systems. Online tutorial, User reference manual, and download site of SiGN-BN are shown in the website (http://sign.hgc.jp/signbn/index.html).

### Cell culture

The human astrocytoma cell line U251 MG (KO) was purchased from JCRB Cell Bank (Tokyo, Japan). U251 MG (KO) cells were cultured in E-MEM containing L-glutamine, phenol red, sodium pyruvate, non-essential amino acids, and 1,500 mg/L sodium bicarbonate (Wako; #055-08975) supplemented with 10% fetal bovine serum (Life Technologies; #10437085) and 100 U/ml penicillin-streptomycin (Life Technologies; #15140122) under 5% CO2 at 37°C. To induce DNA damage, cells were treated with 10-100 nM CLM (MedChem Express; #HY19609) with or without 1-10 μM NU7441 (Wako; #143-09001), a high-potency selective DNA-PK inhibitor. Mouse motor neuron-like cell line, NSC-34 was obtained from Cosmo Bio (#CLU140). The cells were maintained in DMEM (Sigma; # D5796-500ML) with 10% FBS (ATCC, #30-2020) and 100 U/ml Penicillin-Streptomycin (Invitrogen, #15140122) in 5% CO2 at 37 °C. For the treatment of trypsin, the cells were treated with 0.25% Trypsin-EDTA (Invitrogen, # 25200056) for 5 min at 37°C.

### siRNA transfection

For siRNA transfection, U251 MG (KO) cells were plated on non-coated CellCarrier 96-well plates (PerkinElmer; #6005550) or 6-well plates (Corning) with transfection reagents containing the siRNAs (final concentration: 6 nM in culture medium) and Lipofectamine RNAiMAX Transfection Reagent (Thermo Fisher Scientific; #13778030) in Opti-MEMI following the manufacturer’s instructions. The siRNAs used are listed below (large letters: RNA; small letters: DNA). siNC#1 (negative control siRNA in human, mouse, and rat cells), Life Technologies, #4390844 siNC#2 (negative control siRNA in human, mouse and rat cells), Life Technologies, #4390847 siPRKDC#1 (human PRKDC siRNA), Life Technologies, s773 Sense, 5′-GCGUUGGAGUGCUACAACAtt-3′ Antisense, 5′-UGUUGUAGCACUCCAACGCgg-3′ siPRKDC#2 (human PRKDC siRNA), Life Technologies, s774 Sense, 5′-GCGCUUUUCUGGGUGAACUtt-3′ Antisense, 5′-AGUUCACCCAGAAAAGCGCgg-3′ siPRKDC#3 (human PRKDC siRNA), Life Technologies, s775 Sense, 5′-CAAGCGACUUUAUAGCCUUtt-3′ Antisense, 5′-AAGGCUAUAAAGUCGCUUGaa-3′ For the miRNA mimic and siRNA transfection into NSC-34 cells, the cells were plated on the collagen-coated CellCarrier-96 (PerkinElmer; #6005920) or 6 well plate (Corning, #356400) with the transfection reagents containing 60 nM siNC, siFus#1, siFus#4, siFus#5, siTimeless#1, siTimeless#3, miR-125b-5p mimic, miR-124-3p, pre-miR-125b and Lipofectamine RNAiMAX Transfection Reagent (Thermo; #13778030) in Opti-MEMI (Thermo; #31985070) following the manufacturer’s instructions. miR-125b-5p mimic and pre-miR-125b in this study are listed below. miR-125b-5p; hsa-miR-125b-5p mirVana^®^ miRNA mimic (#4464066, MC10148, ABI), pre-miR-125b; hsa-miR-125b-5p Pre-miR™ miRNA Precursor (#AM17100, PM10148, ABI), miR-124-3p; mmu-miR-124-3p mirVana^®^ miRNA mimic (#4464066, MC10691, ABI), siFus#1 (mouse Fus siRNA), Life Technologies, s107706 Sense, 5′-GGAUAAUUCAGACAACAAUtt-3′ Antisense, 5′-AUUGUUGUCUGAAUUAUCCtg-3′ siFus#4 (mouse Fus siRNA), Life Technologies, s107705 Sense, 5′-GAAGUGUCCUAAUCCUACAtt-3′ Antisense, 5′-UGUAGGAUUAGGACACUUCca-3′ siFus#5 (mouse Fus siRNA), Life Technologies, s107707 Sense, 5′-CGACUGGUUUGAUGGUAAAtt-3′ Antisense, 5′-UUUACCAUCAAACCAGUCGat-3′ siTimeless#1 (mouse Timeless siRNA), Life Technologies, s75146 Sense, 5′-AGAUGUAGUGGAAACCAUAtt-3′ Antisense, 5′-UAUGGUUUCCACUACAUCUtg-3′ siTimeless#3 (mouse Timeless siRNA), Life Technologies, s202282 Sense, 5′-GGAAGAUCCGGAAGAGGAAtt-3′ Antisense, 5′-UUCCUCUUCCGGAUCUUCCtg-3′ Quantification of mRNA by quantitative RT-PCR For quantification of human PRKDC mRNA and human β-actin mRNA expression levels, one-step Real Time-PCR analyses were performed with a TaqMan^®^ Gene Expression assay (Life Technologies; human PRKDC, Hs04195439_s1, β-actin, Hs01060665_g1), FastLane Cell Probe Kit (Qiagen; #216413), and ViiA7 Real Time PCR system (Life Technologies) according to the manufacturer’s instructions. For quantification of gene expression levels in NSC-34 cells, total RNAs were extracted using the miRNeasy Mini kit (Qiagen) and quantified by Qubit with the Qubit^®^ RNA HS Assay Kit (Life Technologies). cDNA was reverse-transcribed from these total RNA samples with SuperScript VILO cDNA synthesis Kit (Life Technologies, #11754050). For quantification of mouse Lin28A, Ppp1ca, Dusp6, Fus, Timeless, Prkdc, and β-actin mRNAs, miR-125b-5p and U5 snRNA expression levels, Real Time-PCR analyses were performed with a TaqMan^®^ Gene Expression assay (Life Technologies, Lin28A, Mm00524077_m1; Ppp1ca, Mm00453295_m1; Dusp6, Mm00518185_m1; Fus, Mm00836363_g1; Prkdc, Mm01342967; Timeless, Mm00495610; mouse β-actin, Mm02619580_g1), microRNA RT Kit (miR-125b-5p, TM000449 and RT000449; U6 snRNA, TM001973 and RT001973), Universal PCR Master Mix II (Life technologies, #4440038) and ViiA7 Real Time PCR system (Life Technologies) according to the manufacturer’s instructions.

### Western blotting

U251 MG (KO) cells and NSC-34 cells were collected into 1× SDS Blue loading buffer (NEB; #B7703S) and heat-treated at 95°C for 5 min. Next, genomic DNA in the samples was crushed using the syringe needle (Terumo; #SS-05M2913). Electrophoresis experiments were performed with TGX AnyKD gels, 7.5% gels (Bio-Rad; #4569036), and 12.5% SuperSep Phos-tag gels (Wako; #195-17991) and blotting experiments were performed with a Transblot Turbo blotting system and PVDF PAK MINI (Bio-Rad, Hercules, CA; #1794156). The blotted membranes were blocked with 5% nonfat dry milk (Cell Signaling Technology; #9999S) in Tris-buffered saline containing Tween-20 (TBST) (Cell Signaling Technology; #9997S) for 30 min at room temperature. The membranes were then incubated with mouse monoclonal anti-phospho-Histone H2A.X (Ser139) (Millipore; #05-636), rabbit polyclonal anti-DNA-PK (Abcam; #ab70250), rabbit polyclonal anti-phospho-DNA-PK (Ser2056) (Abcam; #ab18192), rabbit polyclonal anti-TDP-43 (Proteintech; #1078-2-AP), mouse monoclonal anti-FUS (Santa Cruz Biotechnology; #SC-47711), and anti-β-actin (Novus Biologicals; #NB600-532) primary antibodies in the blocking buffer overnight at 4°C. After three washes with TBST, the membranes were incubated with horseradish peroxidase-conjugated anti-mouse IgG (Cell Signaling Technology; #7076P2) or horseradish peroxidase-conjugated anti-rabbit IgG (Cell Signaling Technology; #7074P2) secondary antibodies in TBST for 30 min at room temperature. Luminescence signals on the membranes were detected by an image reader (LAS-4000 system; Fujifilm) with SignalFire ECL Reagent (Cell Signaling Technology; #6883S). The obtained images were processed by Multi Gauge version 3.1 software equipped with the LAS-4000 system.

### Immunofluorescence microscopy

U251 MG (KO) cells were cultured on non-coated CellCarrier 96-well plates. NSC-34 cells were cultured on collagen-coated CellCarrier 96-well plates. After washing with phosphate-buffered saline (PBS) (Wako; #166-23555), the cells were fixed with 4% paraformaldehyde/phosphate buffer solution (Wako; #163-20145) for 15-30 min on ice and permeabilized three times with 0.1% Triton X-100 in high-salt buffer (500 mM NaCl, 1 mM NaH2PO4.2H2O, 9 mM Na2HPO4, 0.1% Tween-20) for 10 min at room temperature. The cells were then treated with 1% bovine serum albumin (Sigma; #A7030-100G) for 30 min at room temperature and incubated with mouse monoclonal anti-phospho-Histone H2A.X (Ser139) (Millipore; #05-636), mouse monoclonal anti-FUS (Santa Cruz Biotechnology; #sc-47711; diluted 1:250), rabbit polyclonal anti-G3BP1 (Bethyl Laboratories; #A302-033A; diluted 1:250), rabbit polyclonal anti-TDP-43 (Proteintech; #1078-2-AP; diluted 1:250), and rabbit polyclonal anti-hnRNP A1 (Cell Signaling Technology; #8443S; diluted 1:250) primary antibodies in low-salt buffer (0.05% Tween-20 in PBS) overnight at 4°C. After three 5-min washes with high-salt buffer at room temperature, the cells were incubated with Alexa Fluor 488 Goat Anti-Rabbit IgG (H+L) (Invitrogen; #A11034) and Alexa Fluor 594 Goat Anti-Mouse IgG (H+L) (Invitrogen; #A11032) secondary antibodies in low-salt buffer for 30 min at room temperature. After three 5-min washes with high-salt buffer at room temperature, the immunofluorescence signals were observed under a fluorescence microscope (Keyence; BZ-X710) equipped with ×20 (Nikon; PlanApo λ, NA = 0.75) or ×40 (Nikon; PlanApo λ, NA = 0.95) objective lens, and images were obtained with the equipped software (BZ-X Viewer and Analyzer).

### RNA-Seq and GO term analyses

Total RNAs were extracted from the NSC-34 cells using the miRNeasy Mini kit (QIAgen) and quantified by Qubit with the Qubit^®^ RNA HS Assay Kit (Life Technologies). RNA-Seq service of strand-specific RNA library preparation with poly A selection were provided with Illumina HiSeq, 2×150bp by GeneWiz (South Plainfield, NJ). Obtained FASTQ files were processed through Trimmomatic, PRINESQ, STAR, featureCounts and edgeR algorithms. For GO analyses with DEGs between the cells in the control conditions (non-siRNA, siNC#1, and siNC#2) and overexpressed with miRNAs (miR-125b-5p mimic, pre-miR-125b precursors and miR-124-3p mimic), down-regulated DEGs (logFC < 0.5, P-Value < 0.05) in the cells treated with miR-125b-5p mimic, pre-miR-125b and miR-124-3p mimic were applied to GO analysis with DAVID Bioinformatics Resources 6.8. RNA-seq dataset of NSC-34 cells treated with siRNA (-), siNC#1, siNC#2, siFus#1, siFus#4, siFus#5, miR-125b-5p mimic, miR-124-3p mimic and pre-miR-125b have been deposited with GEO under accession number GSE125655.

## DATA ACCESS

The data discussed in this publication have been deposited in NCBI’s Gene Expression Omnibus (GEO) and are accessible through GEO Series accession number GSE118336 and GSE125655.To access the dataset GSE118336, please visit the website as shown below and enter secure token. The following secure token has been created to allow review of record GSE118336 while it remains in private status:

secure token: upybwgagrvojtit

https://urldefense.proofpoint.com/v2/url?u=https-3A_www.ncbi.nlm.nih.gov_geo_query_acc.cgi-3Facc-3DGSE118336&d=DwIEAg&c=FyTjmTD2fsLzxJqwPQvEZg&r=CyqwpOWecMoBGI-vBNGIOgVeqAV1EuBrs7mObQ1P9AU&m=s6zuRU289ga0eWZ4ZbU3LAhZPiQtDhOcrPuBC9udM&s=X2AJQEo-u-kk5pBV3

To access the dataset GSE125655, please visit the website as shown below and enter secure token. The following secure token has been created to allow review of record GSE125655 while it remains in private status:

secure token: klyxygeqvlkzdgz

https://urldefense.proofpoint.com/v2/url?u=https-3A_www.ncbi.nlm.nih.gov_geo_query_acc.cgi-3Facc-3DGSE125655&d=DwIEAg&c=FyTimTD2fsLzxJqwPQvEZg&r=CyqwpOWecMoBGI-vBNGIOgVeqAV1EuBrs7mObQ1P9AU&m=4icW7h0yYTXadsuzSuy6e5DmlLNd-N7m6iDZF3r5K4&s=LCtgdUCdUutaj-TGN

## RESULTS

### Identification of PRKDC as a network “hub gene” by iBRN

To identify the key pathways and genes involved in the molecular aetiology of FUS-ALS, we performed transcriptome analysis of human iPSC-derived MNs harboring mutated FUS^H517D^ and FUS^WT^ (Table 1). In this study, we used three types of iPSCs derived motor neurons, control from iPSCs (409B2 line) as wild type, fALS patient-derived iPSCs as previously reported (Ichiyanagi et al., 2016) as Hetero and homozygous for the FUS^H517D/H517D^ as Homo. All iPSC clones were differentiated into MNs and maintained in the differentiation medium for 2 or 4 weeks as reported previously (Fujimori et al., 2018). Next, we generated large amount of MNs expressing pan-neuron and MN marker (Supplemental Fig. S1A) and performed gene expression profiling. There were no samples that required exclusion for quality problems (Supplemental Fig. S2A). MNs derived from WT (MN-WT) and Hetero clones (MN-Hetero) were differently clustered from Homo clones (MN-Homo) on PCA1 (Supplemental Fig. S2B). MN-Hetero and Homo were separately clustered from MN-WT on PCA3. On PCA2, MNs differentiated from EKA03 clones were differently clustered from 201B7 and 409B2 clones. These results would reflect 201B7 and 409B2 clones were established from the same person andEKA03 was established from another person. Next, the gene expression levels of neuron and MN markers were investigated (Supplemental Fig. S1B-G). ISL1 was comparatively up-regulated in MN-Hetero, while the other marker gene expression did not differ much among MN-WT, MN-Hetero and MN-Homo, therefore, the FUS^H517D^ mutation had no major impact on MN differentiation and our large-scale transcriptome data would have the potential to identify key genes and pathways. Numbers of significantly differential expressed genes (DEGs) between the 2 and 4 weeks groups in each genotype comprised approximately 0.5% of the whole transcripts (Supplemental Fig. S2C). Number of DEGs between MN-WT and MN-Hetero comprised approximately 1.6% of the whole transcripts (Supplemental S2D). Meanwhile, number of DEGs between MN-WT versus MN-Homo, and MN-Hetero versus MN-Homo exceeded 3.5% of the whole transcripts. The common 219 DEGs of 4 comparisons, Hetero_2ws versus WT_2ws, Hetero_4ws versus WT_4ws, Homo_2ws versus WT_2ws, and Homo_4ws versus WT_4ws were used to Gene Ontology (GO) term analyses, “Cell proliferation” and “DNA repair”-related terms were identified in the top term list (Supplemental Fig. S2E). Actually, large portion of the down-regulated DEGs associating with “DNA repair” and “DNA replication” terms were shared by both terms. A GO term analysis of the down-regulated 512 DEGs in MN-Hetero relative to MN-WT was performed and “DNA repair”-related GO terms were identified in the top list (Supplemental Fig. S2F). Similarly, DDR-related pathways were uniformly down-regulated in MN-Hetero and MN-Homo from pathway analyses (Supplemental Fig. S3). Numbers of the DEGs in the top pathways were augmented in the comparison between Homo versus WT groups (Supplemental Fig. S3C). Collectively, these results indicated that FUS^H517D^ mutation would cause uniform down-regulation of DDR-related genes expression.

**Table 1.**
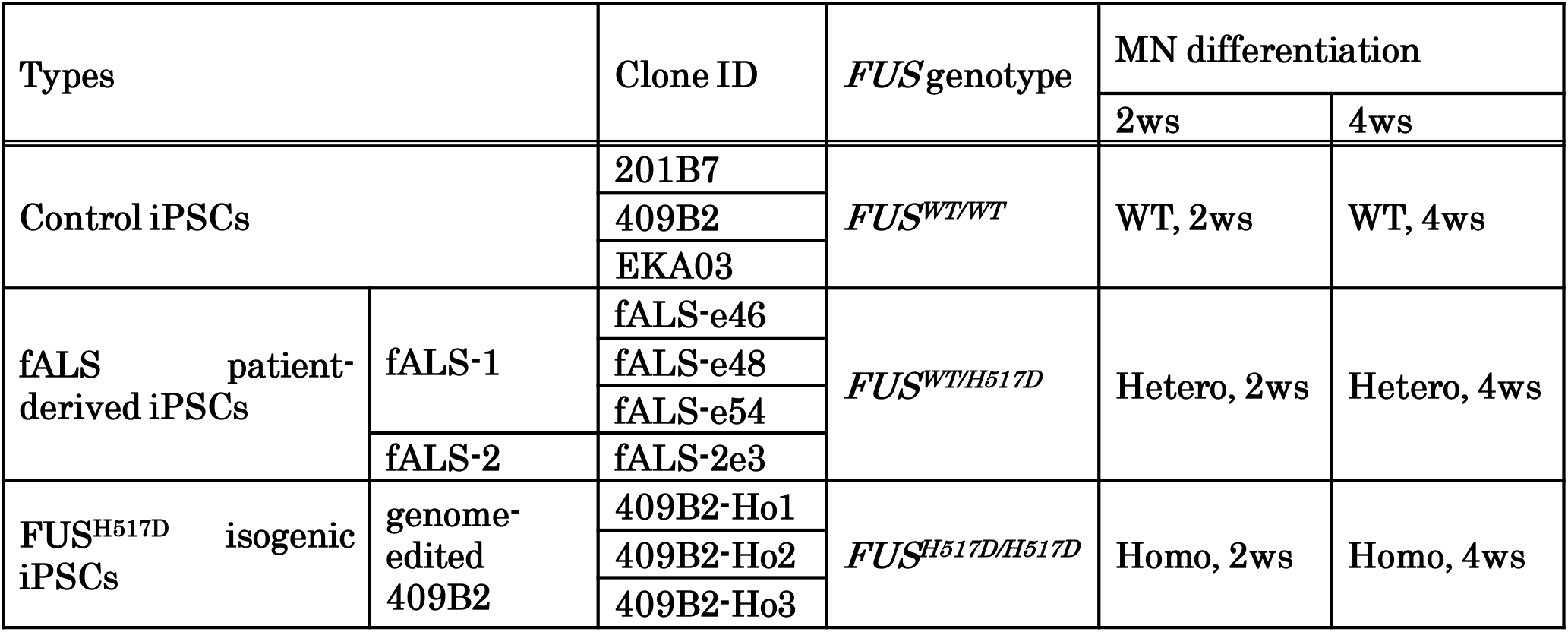
Sample information for iPSC-derived MNs in this study. Sample information for the iPSC-derived MNs used in the large-scale transcriptome analysis with the GeneChip HTA2.0 and nCounter miRNA analysis system is shown. Total RNA was purified from iPSC- derived MNs differentiated for 2 weeks (shown as MN differentiation, 2ws) and 4 weeks (shown as MN differentiation, 4ws) from MPCs. 201B7, 409B2, and EKA03 iPSC lines were used as control cells harboring FUS^WT/WT^. Familial ALS (fALS)-1 and fALS-2 were independent patients and iPSCs derived from these patients and genome-edited 409B2 iPSCs were used as cells harboring FUS^WT/H517D^and FUS^H517D/H517D^, respectively (Ichiyanagi et al., 2016). 10 iPSC clones were triplicatedly differentiated into MNs for 2 or 4 weeks and 60 total RNA samples were purified.

| Types |  | Clone ID | *FUS* genotype | MN differentiation |  |
| --- | --- | --- | --- | --- | --- |
|  |  |  |  | 2ws | 4ws |
| Control iPSCs |  | 201B7 | *FUS<sup>WT/WT</sup>* | WT, 2ws | WT, 4ws |
|  |  | 409B2 |  |  |  |
|  |  | EKA03 |  |  |  |
| fALS patient-derived iPSCs | fALS-1 | fALS-e46 | *FUS<sup>WT/H517D</sup>* | Hetero, 2ws | Hetero, 4ws |
|  |  | fALS-e48 |  |  |  |
|  |  | fALS-e54 |  |  |  |
|  | fALS-2 | fALS-2e3 |  |  |  |
| *FUS<sup>H517D</sup>* isogenic iPSCs | genome-edited 409B2 | 409B2-Ho1 | *FUS<sup>H517D/H517D</sup>* | Homo, 2ws | Homo, 4ws |
|  |  | 409B2-Ho2 |  |  |  |
|  |  | 409B2-Ho3 |  |  |  |

iBRN analysis can identify “hub genes”, which are expected to play key roles in a gene network of iPSC-derived cells (Fig. 1A). In the first analysis, ALS-related gene network was calculated with 754 genes related to ALS-related pathways (including ALS causal genes, mitochondria, pre-mRNA and miRNA processing, p53 and RB signalling, DDR, prion disease, and kinesin motor proteins) and the top hub genes were identified (Fig. 1B and Supplemental Table S1). In the second analysis, 1,542 genes encoding RBPs (Gerstberger et al., 2014) were focused upon to estimate the RBP gene network, because ALS is known as an RNA metabolic disease (Fujimori et al., 2018, Barmada, 2015, Gama-Carvalho et al., 2017, Taylor et al., 2016, Akiyama et al., 2019) and the 900 RBP genes which expresses well in our samples were used for iBRN (Fig. 1C and Supplemental Table S1). PRKDC (alternative name, DNA-dependent protein kinase, catalytic subunit, DNA-PKcs), encoding a responsible kinase for phosphorylation of FUS during DNA damage stress and negative regulator of FUS liquid droplet and fibril formation in vitro (Han et al., 2012, Deng et al., 2014, Monahan et al., 2017, Monahan et al., 2017, Murray et al., 2017) was identified as a top hub gene in both gene networks (Fig. 1B and C, Supplemental Table S1). Surprisingly, PRKDC expression levels clearly showed the second best correlation with FUS expression levels in FUS^WT^ MNs and FUS^H517D^ MNs (Fig. 1D). PRKDC expression levels in the Hetero_2ws and Homo_2ws groups were down-regulated to two-thirds and one-third, respectively, compared with the WT_2ws group (Fig. 1E). FUS expression levels were similarly down-regulated in MN-Hetero and MN-Homo compared with MN-WT (Fig. 1F) and the number of FUS^H517D^ mutations was inversely correlated with PRKDC and FUS expression levels (Fig. 1E and F).

**Figure 1.**
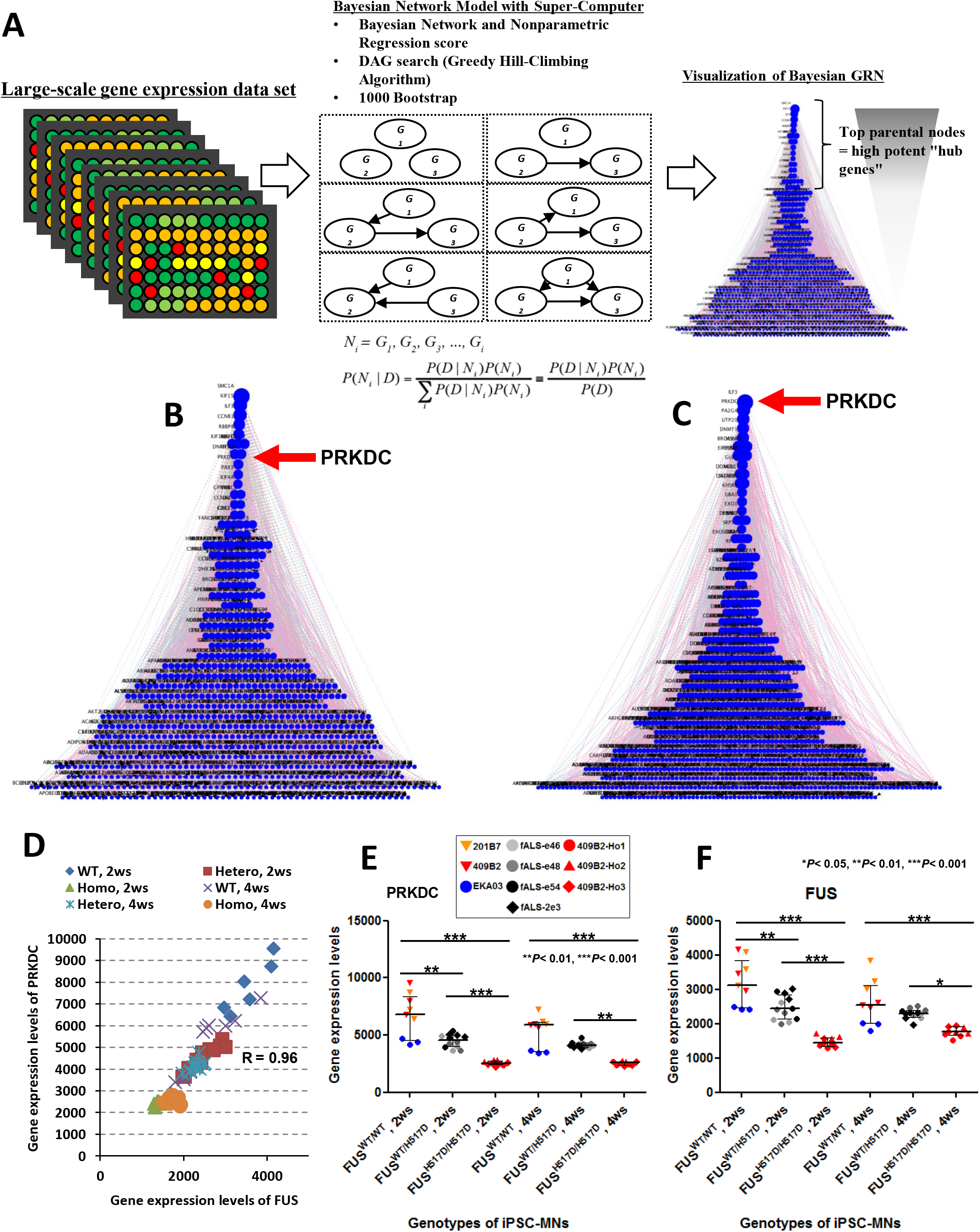
Identification of PRKDC as a “hub gene” by iBRN. (A) Workflow of iBRN with large transcriptome dataset. The Bayesian network is a directed acyclic graph (DAG) with probabilistic transition on its edges. When gene expression data (D) is given, a gene regulatory network can be estimated by the DAG search results. In this case (i=3), we have 6 structures of network that means 25 DAGs. If we have 30 genes (i=30), the number of DAGs exceeds 2.71 × 10158, therefore, the estimation requires a huge amount of computational resources. The regulatory networks were visualized with nodes and edges which represent for genes and relationships, respectively. All edges have direction between nodes and the parental nodes possessing many children nodes were laid out in the top of the network. Top parental nodes are recognized as “hub genes” in the regulatory networks. The color of edges indicates the relation types; pink = up regulation, blue = down regulation and gray = unknown relation. (B) The 754 genes corresponding to ALS-related pathways were selected for iBRN. The gene expression data of these genes in 60 samples were used for the analysis. The calculated network is shown in the Sugiyama layout. The genes in the upper part of the triangle are strong “hub genes”. Circle size indicates the number of “children” genes of each gene. The red arrow indicates the position of the PRKDC gene. Top 20 hub genes are shown in the left table of Supplemental Table S1. (C) The 900 genes encoding RBPs were selected for iBRN. The red arrow indicates the position of the PRKDC gene. Top 20 hub genes are shown in the right table of Supplemental Table S1. (D) PRKDC showed the second-best correlation with FUS in expression (R= 0.96). The gene expression levels of PRKDC (E) and FUS (F) in the WT_2ws (n = 9), Hetero_2ws (n =12), Homo_2ws (n = 12), WT_4ws (n = 9), Hetero_4ws (n = 12), and Homo_4ws (n = 12) group are shown graphically. Values represent mean ± SD and the star indicates *P< 0.05, **P< 0.01, or ***P<0.001 (Tukey’s multiple comparison test).

### PRKDC functions in regulating localization of FUS protein under DNA damage stress

As PRKDC was identified to be a potential “hub gene” in the regulatory networks by iBRN, we validated the functional interactions among FUS, PRKDC, and DNA damage stress in the culture cells. In normal conditions, FUS was localized at the nucleus in U251 MG (KO) cells (Fig. 2A). Upon treatment with a DNA double-strand break inducer calicheamicin (CLM), the accumulation of DNA damage (Supplemental Fig. S4A) and G3BP1-positive cytosolic SG formations (Fig. 2A) were observed. Meanwhile, FUS was still localized in the nucleus (Fig. 2A). The treatment with a DNA-PK-specific inhibitor (NU7441) also showed that FUS was still localized in the nucleus (Fig. 2A). Interestingly, after co-treatment with both NU7441 and CLM, cytosolic mislocalization of FUS was observed in G3BP1 positive SGs (Fig. 2A and B). Importantly, the FUS protein band was shifted upward in a CLM-dependent manner and this upshift was totally suppressed by NU7441 treatment, suggesting the protein phosphorylation of FUS (Fig. 2C). Western blotting analysis with a Phos-tag SDS-PAGE gel, an electrophoresis technique capable of separating phosphorylated and non-phosphorylated proteins based on phosphorylation levels, further confirmed dramatic migration of the FUS protein with the treatment of CLM (Fig. 2D). These results indicated that phosphorylation of FUS depends on DNA-PK activity. In addition, after treatment with three independent siRNAs against PRKDC mRNA, suppression of PRKDC mRNA and protein showed cytosolic mis-localization of FUS as a common feature of the cells (Fig. 2E-G). Collectively, down-regulation of DDR-related gene, PRKDC had impact on the gene network through the intracellular FUS mis-localization. However, PRKDC specifically play a role in localization of FUS, not to the other ALS causal RBPs, such as TDP-43 and hnRNPA1 (Izumi et al., 2015, Kim et al., 2013) (Supplemental Fig. S4B_D). These results suggested that PRKDC could specifically act as a guardian against FUS mis-localization at cytosolic SGs during DNA damage stress and maintain FUS nuclear localization (Supplemental Fig. S4E).

**Figure 2.**
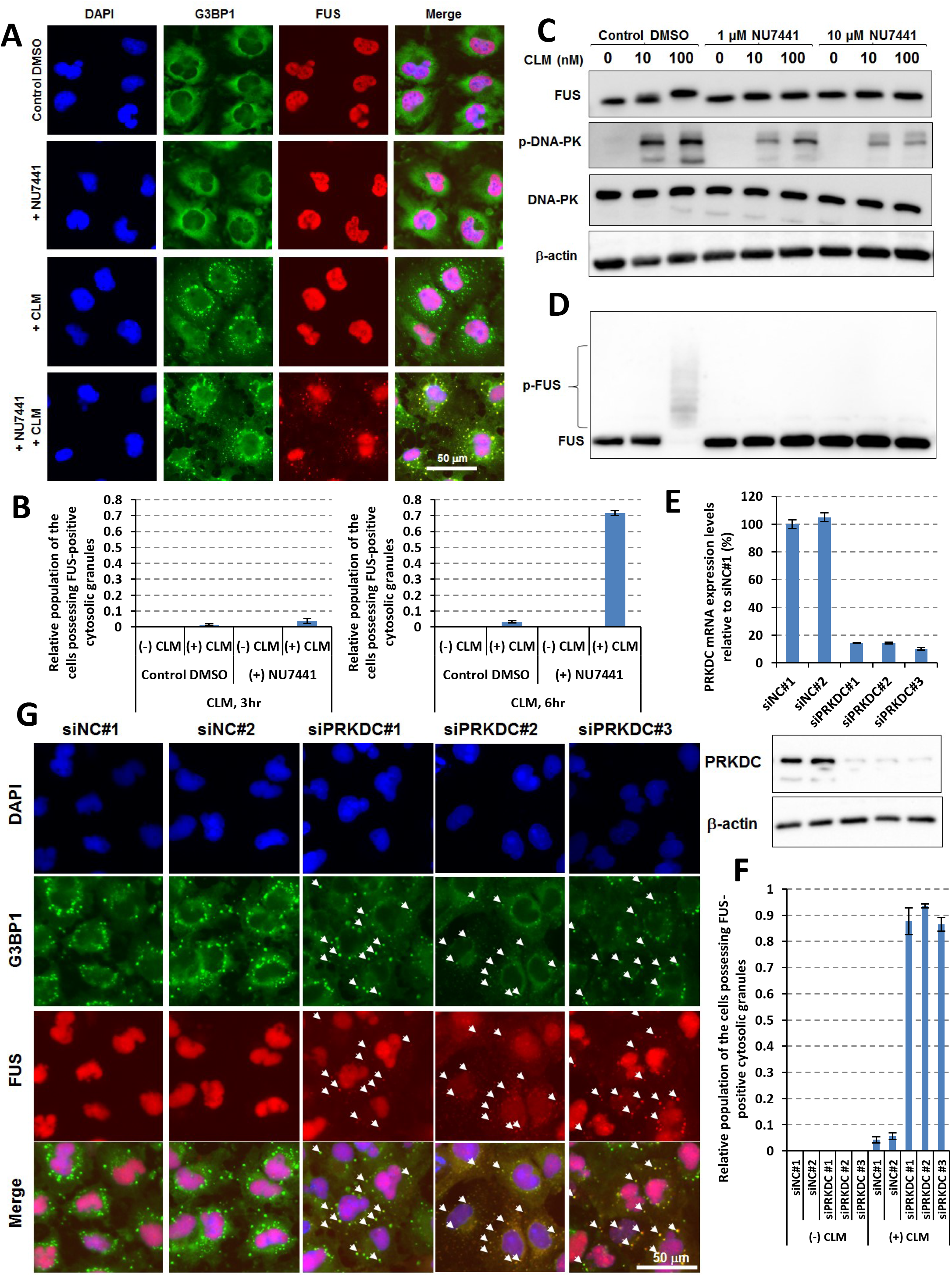
PRKDC functions in regulating localization of FUS protein under DNA damage stress. (A) U251 MG (KO) cells were pre-treated with control dimethyl sulfoxide (DMSO) or 1 μM NU7441 and then treated with control DMSO or 100 nM CLM for 6 hr. Immunofluorescence imaging was performed with DAPI, anti-FUS, and anti-G3BP1 antibodies. (B) Four fields of cells for each condition were imaged and the populations of cells possessing FUS-positive cytosolic SGs were counted. Relative populations to the total cell number (monitored with DAPI) after 3 hr (shown in left graph) or 6 hr (shown in right graph) of stimulation with CLM were calculated. (C) U251 MG (KO) cells were pre-treated with control DMSO or 1-10 μM NU7441 for 3 hr and then treated with control DMSO or 10-100 nM CLM for 3 hr. Western blotting analysis was performed with anti-FUS, anti-DNA-PK, anti-phospho-S2056-DNA-PK and anti-β-actin antibodies. (D) A Phos-tag gel shift assay was performed with an anti-FUS antibody to detect phosphorylated FUS (p-FUS). (E) U251 MG (KO) cells were pre-treated with PRKDC siRNA for 2 days. The PRKDC mRNA expression levels relative to β-actin mRNA expression levels were calculated. Cells treated with negative control siRNA #1 (siNC#1) are shown as 100%. For western blotting, U251 MG (KO) cells were pre-treated with PRKDC siRNA for 3 days. The PRKDC protein and β-actin proteins as the loading control were analysed by the specific antibodies. U251 MG (KO) cells were pre-treated with PRKDC siRNA for 3 days and then treated with 100 nM CLM for 6 hr. Immunofluorescence imaging was performed with DAPI, anti-FUS, and anti-G3BP1 antibodies and four fields of cells in each condition were imaged (G). Arrowheads indicate the cytosolic granules where FUS and G3BP1 proteins were co-localized. The populations of the cells possessing FUS-positive cytosolic granules were counted and the relative populations to the total cell number (monitored with DAPI) were calculated (F).

### Identification of miR-125b-5p as a “hub miRNA”

A miRNA is a small non-coding RNA that functions in negative regulation of multiple genes in expression through direct interactions with the 3′-UTRs of the target mRNAs. It was previously reported that FUS functions in the production of miR-9-5p, a key player in neuronal function (Morlando et al., 2012). Considering the possibility that expression of mutated FUS proteins result in dysregulation of miRNA production including miR-9 and impairment of MNs, we performed a miRNA profiling and iBRN analysis with MN-WT and MN-Homo. A clustering analysis with the top 179 expressed miRNAs showed separate groups between the MN-WT and MN-Homo (Fig. 3A). Indeed, miR-9-5p expression levels were down-regulated in the MN-Homo compared with the MN-WT (Fig. 3B). These results supported the notion that expression of mutated FUS proteins results in dysregulation of miR-9-5p production, as reported previously (Morlando et al., 2012). Furthermore, we found that miR-125b-5p was dramatically up-regulated in the MN-Homo compared with the MN-WT (Fig. 3C). Taken together, these results indicated that the mutated FUS gene would have both positive and negative impacts on dysregulation of miRNA production. Next, the gene networks including miRNAs including 110 miRNAs, which were abundantly expressed in the samples and genes shown in Fig. 1B and C, identified miR-125b-5p as “hub miRNA” in both networks (Fig. 3D and Supplemental Table S2). Alternatively, miR-205-5p was another miRNA identified as “hub miRNA” in both networks and down-regulated in the Homo group in comparison with the WT group (Fig. 3E).

**Figure 3.**
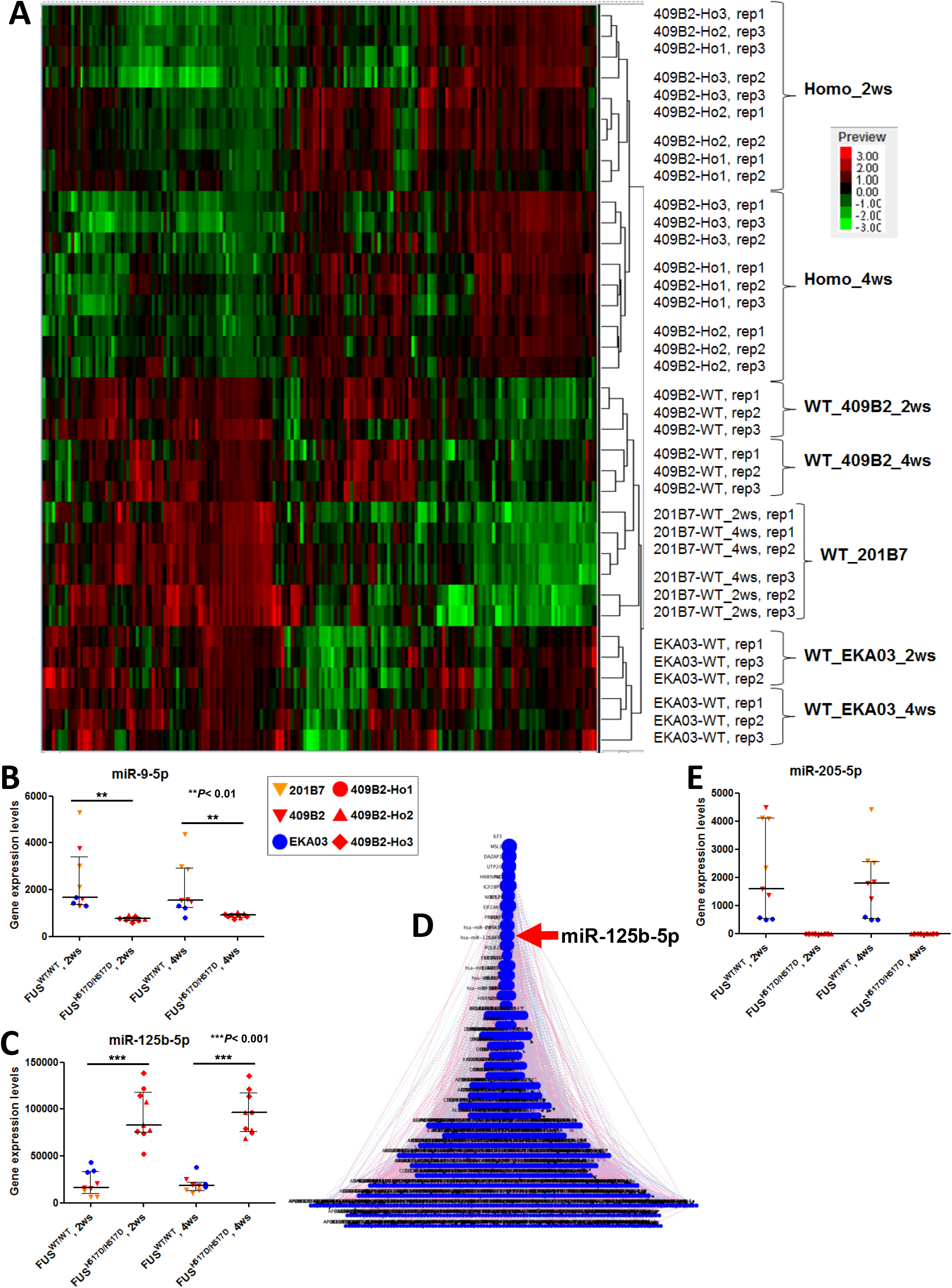
Identification of miR-125b-5p as a “hub miRNA” by iBRN. (A) clustering analysis with the top 179 expressed miRNAs (signal values > 100) was performed. (B) The hsa-miR-9-5p expression levels were down-regulated in MN-Homo compared with MN-WT. Values represent mean ± SD (unpaired Student’s t test). (C) The hsa-miR-125b-5p expression levels were up-regulated in MN-Homo relative to MN-WT. Values represent mean ± SD (unpaired Student’s t test). (D) The 900 RBP genes selected in Fig. 1B and 110 miRNAs were used for iBRN. The red arrow indicates the position of miR-125b-5p. Top 20 hub genes are shown in the right table of Supplemental Table S2. (E) The hsa-miR-205-5p expression levels were down-regulated in MN-Homo relative to MN-WT.

### Correlation analyses of miR-125b-5p expression

Mature forms of miR-125b-5p are processed from pri-miR-125b-1 and pri-miR-125b-2, which are encoded in the introns of MIR100HG and MIR99AHG genes on chromosome 11 and 21, respectively. Pri-miR-125b-1 and MIR100HG clearly showed good correlation to mature miR-125b-5p (Fig. 4A and B) and were up-regulated in MN-Homo (Fig. 4C and D), but pri-miR-125a and pri-miR-125b-2 did not (Supplemental Fig. S5A and B). MIR100HG gene encodes miR-100-5p and let-7a and both miRNA expression levels were similarly up-regulated in MN-Homo (Fig. 4E and F). These results suggested that mutated FUS^H517D^ would cause transcriptional activation of MIR100HG gene and following accumulation of miR-125b-5p mainly processed from pri-miR-125b-1. LIN28A, Dual-specific phosphatase 6 (DUSP6), and protein phosphatase 1 catalytic subunit alpha isozyme (PPP1CA) mRNAs were reported to be targets of miR-125b-5p (Banzhaf-Strathmann et al., 2014; Chaudhuri et al., 2012), and their gene expression levels were down regulated in FUS^H517D^ MNs (Fig. 4G_I) and showed negative correlation to miR-125b-5p in MN-WT and MN-Homo (Fig. 4J_L).

**Figure 4.**
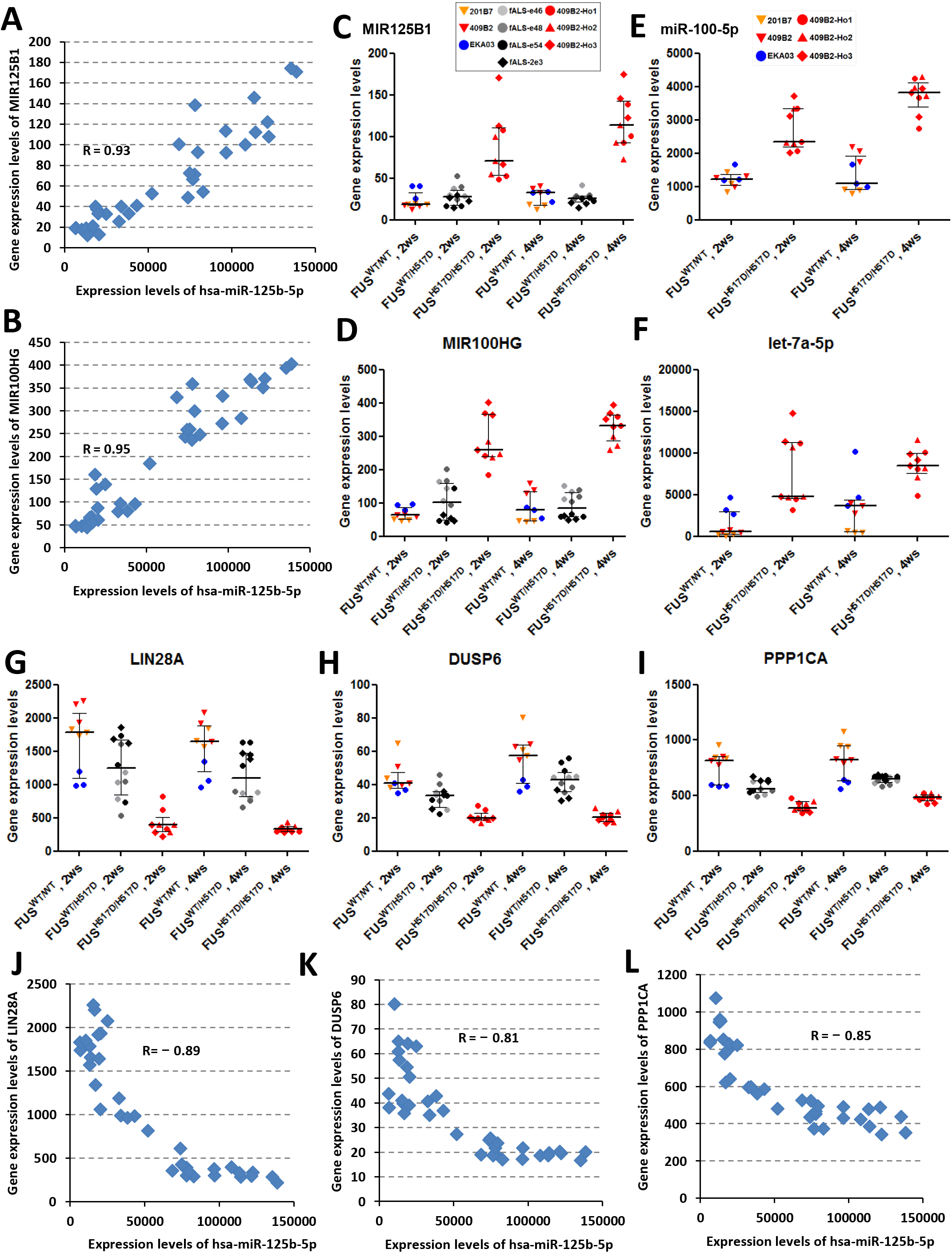
Correlation analyses of miR-125b-5p expression. (A) A positive correlation was observed between pri-miR-125b1 gene expression and hsa-miR-125b-5p expression (R = 0.93). (B) A positive correlation was observed between MIR100HG gene expression and hsa-miR-125b-5p expression (R = 0.95). The gene expression levels of pri-miR125b1 (C) and MIR100HG (D) in the WT_2ws (n = 9), Hetero_2ws (n = 12), Homo_2ws (n = 12), WT_4ws (n = 9), Hetero_4ws (n = 12), and Homo_4ws (n = 12) group are shown graphically. Values represent mean ± SD. The hsa-miR-100-5p (E) and hsa-let-7a-5p (F) expression levels were up-regulated in MN-Homo relative to MN-WT. Values represent mean ± SD). The LIN28A (G), DUSP6 (H) and PPP1CA (I) gene expression levels were down-regulated in the FUS^H517D^ MNs relative to the FUS^WT^ MNs. Values represent mean ± SD. Negative correlations of hsa-miR-125b-5p with LIN28A (J), DUSP6 (K) and PPP1CA (L) expression levels were shown.

### Up-regulation of miR-125b-5p causes DNA damage

Next, we assessed these mRNAs as potential targets of miR-125b-5p in cells (Banzhaf-Strathmann et al., 2014, Chaudhuri et al., 2012, Chaudhuri et al., 2012). The gene expression levels of Lin28a, Dusp6 and Ppp1ca were down-regulated by the treatment with miR-125b-5p mimic and pre-miR-125b in NSC-34 cells (Fig. 5A_C). Surprisingly, Fus expression levels were down-regulated as well (Fig. 5D) and showed clearly negative correlation to miR-125b-5p in MN-WT and MN-Homo (Fig. 5E). On the other hand, other ALS causal genes, TARDBP and C9ORF72 did not show negative correlations to miR-125b-5p (Supplemental Fig. S5C and D). miR-125b-5p is abundant miRNA in human spinal cord and was up-regulated in the lumbar spinal cord of SOD1-G93A mice, reported previously (Ludwig et al., 2016, Parisi et al., 2016). In addition, direct interactions between FUS mRNA and miR-125b-5p have been experimentally supported by HITS-CLIP (Hafner et al., 2010, Balakrishnan et al., 2014, Pillai et al., 2014, Karagkouni et al., 2018). Collectively, up-regulation of miR-125b-5p might function as “hub miRNA” in the acceleration of FUS dysregulation and have impact on FUS-ALS gene networks (Fig. 5F). To better understand miR-125b-5p target mRNA molecules, we performed RNA-seq analyses of NSC-34 cells transiently transfected with miR-125b-5p mimic and pre-miR-125b. Surprisingly, DDR-related genes were down-regulated (Supplemental Fig. S6A). As down-regulated DEGs associating with “Cellular response to DNA damage stimulus” and “DNA repair” GO terms, 61 and 42 genes were identified (Supplemental Table S4). Large portion of these DEGs showed lower expression levels in MN-Hetero and MN-Homo than MN-WT (Supplemental Table S5). Individually, DNA repair-related genes, poly(ADP-ribose) polymerase 1 (Parp1) and Timeless were down-regulated (Supplemental Fig. S6B) (Dash et al., 2017). Numbers of the down-regulated DEGs were augmented in MN-Homo. Introduction of miR-125b-5p specifically down-regulated genes associated with DDR pathway genes (Supplemental Fig. S6C_E) and promoted the accumulation of DNA damage (Fig. 5G and H). Taken together, up-regulation of miR-125b-5p would cause down-regulation of DDR-related genes found in iPSC-derived MNs harboring FUS^H517D^ mutation.

**Figure 5.**
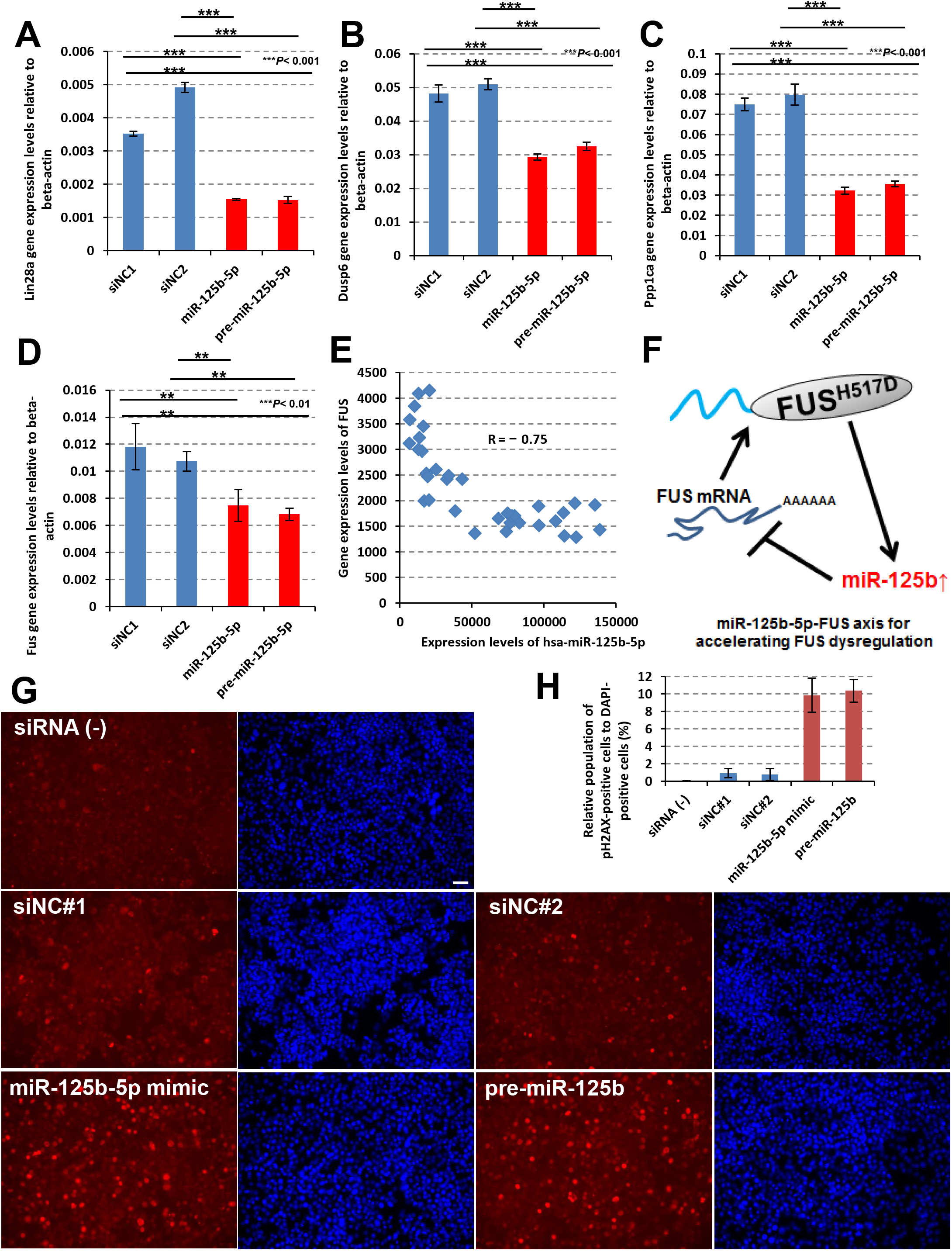
Up-regulation of miR-125b-5p causes DNA damage. Lin28a (A), Dusp6 (B), Ppp1ca (C) and FUS (D) mRNA expression levels were down-regulated by introduction of miR-125b-5p and pre-miR-125b. Values represent mean ± SD (unpaired Student’s t test). (E) A negative correlation was observed between FUS gene expression and hsa-miR-125b-5p expression (R = _0.75). (F) Possible mechanism of miR-125b-5p function in the acceleration of FUS dysregulation as a hub miRNA. (G) NSC-34 cells were transiently transfected with the negative control siRNAs (siNC#1 and siNC#2), miR-125b-5p mimic and pre-miR-125b and after 72 hours from transfection, the cells were fixed to visualize the accumulation of DNA damage by anti-phospho-H2AX antibody (shown in red) and DAPI (shown in blue). A bar in the picture indicates 50 μm. (H) Relative population of the cells possessing phospho-H2AX-positive nucleus were calculated to DAPI-positive cells. Values represent mean ± SD.

### Timeless is a key down-stream regulator of miR-125b-5p

To explore hub genes further, iBRN was performed again with additional Alzheimer’s disease (AD)-associated genes and circadian rhythm-related genes because identified hub molecules, PRKDC and miR-125b-5p were associated with AD pathology and circadian rhythm (Banzhaf-Strathmann et al., 2014, Davydov et al., 2003, Kanungo, 2013, Kanungo, 2013, Cardinale et al., 2012, Ma et al., 2017, Gao et al., 2013, Musiek and Holtzman, 2016). TIMELESS and SST were identified as newly added “hub genes” (Fig. 6A and B, Supplemental Table S3). SST gene expression levels were dramatically up-regulated in MN-Homo (Fig. 6C). TIMELESS expression levels in MN-Hetero and MN-Homo were down-regulated to one-second and one-fourth, respectively, compared with MN-WT (Fig. 6D). TIMELESS expression levels clearly showed negative correlation with miR-125b-5p and Timeless expression levels were down-regulated by the treatment with miR-125b-5p mimic and pre-miR-125b in NSC-34 cells (Fig. 6E and F, Supplemental Fig. S6B and E). These results suggested that TIMELESS would be a target mRNA of miR-125b-5p and form “hub gene-hub miRNA” network. Surprisingly, knocking-down Timeless mRNA levels promoted the accumulation of DNA damage (Fig. 6G_I), although knocking-down Fus expression did not cause the accumulation of DNA damage. Taken together, Timeless is a key regulator of DDR pathway downstream of miR-125b-5p in MNs and reduced expression of Timeless mRNA would have impacted on the gene networks as a network “hub gene”.

**Figure 6.**
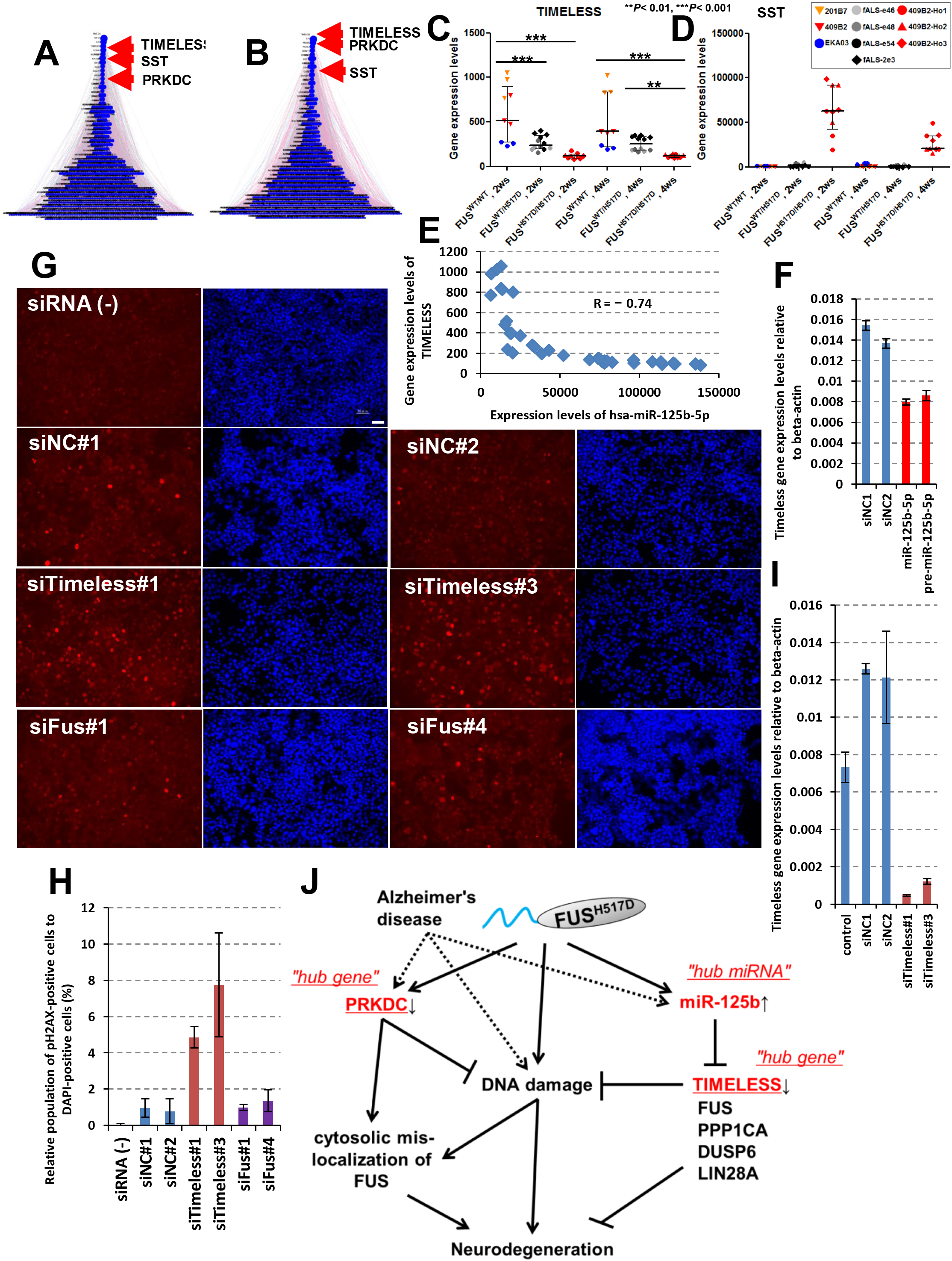
Timeless is a key down-stream regulator of miR-125b-5p. ALS-related pathways with additional genes associated with AD and circadian rhythm were used for iBRN. Top 20 hub genes are shown in the left table of Supplemental Table S3. (B) RBP genes with additional genes associated with AD and circadian rhythm were similarly used for iBRN. Top 20 hub genes are shown in the right table of Supplemental Table S3. (C) TIMELESS gene expression levels are shown graphically. Values represent mean ± SD and the star indicates *P< 0.05, **P< 0.01, or ***P< 0.001 (Tukey’s multiple comparison test). (D) SST gene expression levels were up-regulated in MN-Homo. Values represent mean ± SD. (E) A negative correlation was observed between TIMELESS gene expression and hsa-miR-125b-5p expression (R = _0.74). (F) Timeless mRNA expression levels were down-regulated by the introduction of miR-125b-5p and pre-miR-125b. Values represent mean ± SD (unpaired Student’s t test). (G) Timeless siRNAs and Fus siRNAs were transiently transfected into NSC-34 cells and the accumulation of DNA damage was visualized by anti-phospho-H2AX antibody (shown in red) and DAPI (shown in blue). A bar in the picture indicates 50 μm. (H) Relative population of the cells possessing phospho-H2AX-positive nucleus were calculated to DAPI-positive cells. Values represent mean ± SD. (I) Timeless mRNA expression levels. Values represent mean ± SD. (J) Possible model of hub molecules in FUS-ALS gene network. miR-125b-5p was up-regulated in FUS^H517D^ MN and down-regulates the genes associating with DDR-related pathways which included FUS and TIMELESS. Meanwhile, genes in DNA damage stress response pathway including PRKDC and TIMELESS were uniformly down-regulated in expression and DNA damage under impaired DNA-PK activity condition could promote cytosolic FUS mis-localization to SG

In conclusion, we applied both regular bioinformatics and iBRN to large-scale transcriptome based on iPSC-derived FUS^WT^ and FUS^H517D^ MNs and miR-125b-5p, TIMELESS, and PRKDC were identified as network hub molecules (Fig. 6J). miR-125b-5p was up-regulated in FUS^H517D^ MNs (Fig. 3) and negatively regulates DDR pathway-related genes including FUS and TIMELESS (Supplemental, Fig. S6). In addition, introduction of miR-125b-5p and knocking-down Timeless expression caused the accumulation of DNA damage (Fig. 5G and H, Fig. 6G_I). Meanwhile, PRKDC were down-regulated in FUS^H517D^ MNs and DNA damage under impaired DNA-PK activity condition promoted cytosolic FUS mis-localization to SGs (Supplemental Fig. S4E). Collectively, our strategy using human iPSC model would provide the first compelling evidence to elucidate FUS-ALS molecular aetiology through identification of key hub molecules and their functional analyses.

## DISCUSSION

In the present study, we took biological and computational approaches with large-scale transcriptome based on FUS^WT^ and FUS^H517D^ MNs to uncover molecular aetiology of FUS-ALS. iBRN identified miR-125b-5p, TIMELESS and PRKDC as hub molecules which were strongly affecting the FUS-ALS regulatory gene networks (Fig. 1B, 1C, 3D, 6A and 6B). Traditional pathway analyses of DEGs also found genes associated with DDR pathway including FUS, PRKDC and TIMELESS, which were uniformly down-regulated in FUS^H517D^ MN (Fig. 1E, 1F, 6C, Supplemental Fig. S2 and S3), suggesting that DDR pathway would be key event in the regulatory network. Furthermore, we validated a direct link among hub molecules from a point of view of DNA damage stress with cellular models and provided the evidences for the distinctive roles of these hub molecules in the cytosolic mis-localization of FUS (Fig. 2) and DNA damage accumulation (Fig. 5 and 6). To our knowledge, this study provides the first use that gene network estimation identifies the hub molecules related to the aetiology of FUS-ALS with human iPSC models.

Among three candidate hub genes, miR-125b-5p was up-regulated in FUS H517D MNs (Figure 3C) and the introduction of miR-125b-5p caused the accumulation of DNA damage (Figure 5G) could be due to down-regulation of the genes associating with DDR-related pathways in addition to Fus, Timeless and Parp1 (Figure S6). In fact, large portion of these down-regulated DEGs associating with DDR-related pathways were lower expressed in MN-Hetero than in MN-WT, and the down-regulation was further augmented in MN-Homo (Table S5), suggesting the relationship of mutant FUS protein dose. fALS patients harboring FUS-NLS mutations often exhibited increased DNA damage in the postmortem motor cortex and increased levels of oxidative DNA damage in the spinal cord of sporadic and fALS patients (Ferrante et al., 1997, Wang et al., 2013, Deng et al., 2014). A recent study using iPSC-derived MNs harboring FUS-NLS mutations also discovered inappropriate DDR, which could be a new therapeutic strategy for ALS as a key upstream event (Naumann et al., 2018). These results support that our regulatory networks based on iPSC-derived MNs would reflect the molecular aetiology of FUS-ALS. Furthermore, up-regulation of miR-125b-5p, we found in iBRN, would be a key influence factor to lead MNs degeneration, mediating DNA damages and down-regulation of its targets.

By iBRN analyses, we identified PRKDC which is a key regulator of DDR as one of network hub genes (Fig. 1B and C). DNA-PK is a well-known kinase that directly phosphorylates FUS at Ser/Thr residues located in the low complexity (LC) domain of the N-terminal region and DNA-damage-induced FUS phosphorylation by DNA-PK led to nuclear export of FUS (Deng et al., 2014). Recently, phosphorylation of Ser/Thr on FUS-LC by DNA-PK negatively impacts on FUS-LC self-assembly and liquid-liquid phase separation of FUS-LC droplets (Han et al., 2012, Monahan et al., 2017, Murray et al., 2017). However, the effect of DNA-damage-induced FUS phosphorylation on cytosolic localization of FUS at SGs in cells has not been well investigated so far. In this study, we observed translocation of endogenous FUS^WT^ proteins from nuclear to cytosolic SGs under DNA damage stress condition with impaired DNA-PK activity by specific DNA-PK inhibitor and PRKDC siRNAs (Fig. 2). In addition, PRKDC specifically regulated FUS localization, not to the other ALS causal RBPs, such as TDP-43 and hnRNPA1 (Supplemental Fig. S4B_D). These results suggested that PRKDC could specifically act as a guardian against FUS mis-localization at cytosolic SGs during DNA damage stress and maintains FUS nuclear localization (Supplemental Fig. S4E). Recently, Rhoads and colleagues reported that translocation of endogenous FUS^WT^ proteins from nuclear to cytosol under CLM-induced DNA damage stress condition were promoted by the treatment with the specific DNA-PK inhibitor NU7441 in H4 neuroglioma cells (Rhoads et al., 2018). In addition, nuclear-to-cytoplasmic mis-localization of FUS^WT^ proteins in human and mouse VCP-mutant ALS models and human sporadic ALS were observed (Tyzack et al., 2019). These observations supported our results, however, the function of PRKDC on localization of endogenous FUS^WT^ proteins seemed like controversial against the results reported previously (Deng et al., 2014, Naumann et al., 2018). Output of the phosphorylated form of endogenous FUS^WT^ proteins on shuttling might be different in the cell culture condition and cell types.

In this study, TIMELESS was also identified as one of hub genes (Fig. 6A and B). Mammalian Timeless have been shown to play an essential role in proper progression of DNA replication, activation of cell-cycle checkpoints, and the establishment of sister chromatid cohesion (Mazzoccoli et al., 2016). Human Timeless forms a tight complex with PARP1 and recruitment of Timeless to DNA damage sites depends on the interaction with PARP1, which could promote DNA repair, in particular homologous recombination repair (Young et al., 2015, Xie et al., 2015). Using iPSC-derived MNs, impairment of PARP-dependent DNA damage response signaling due to mutations in the FUS-NLS promotes cytoplasmic FUS mis-localization which results in neurodegeneration and FUS aggregate formation (Naumann et al., 2018). Indeed, we found that knocking-down Timeless expression levels promoted the accumulation of DNA damage (Fig. 6G_I), thus, down-regulation of TIMELESS gene expression might cause misregulation of DNA repair in *FUS^H517D^* MNs. Furthermore, we observed down-regulation of PARP1 gene expression in MN-Homo and miR-125b-5p down-regulated Parp1 gene expression (Supplemental Fig. S5F, S6B and S6E), however, PARP1 was not identified in the top hub gene lists in spite of the existence in those lists (Supplemental Table S1_3). On the other hands, Timeless has been identified as a top network gene in late-onset AD (Zhang et al., 2013), therefore, our regulatory networks may reflect the molecular aetiology of AD beyond FUS-ALS.

Conclusively, we propose biological and non-biased computational approach, iBRN using large-scale transcriptome of human iPSC models would be very powerful strategy to uncover unknown disease aetiology through identification of key hub molecules.

## Author Contributions

M.N., M.I., A.N., M.Y. and H.O. developed hypothesis and designed the experiments. M.I. performed all experiments using hiPSCs. M.I., T.S., T.A., M.A. and H.O. contributed toward hiPSC culture and established isogenic lines. A.D. performed iBRN analyses. M.N. contributed toward microarray analyses, miRNA profiling and RNA-sequencing. M.N. and O.S. performed the examination of cell assays. M.N., M.I., A.D., K.O., M.Y. and H.O. prepared the manuscript and input from all the authors. All authors read and approved the final manuscript.

## Acknowledgments

We thank Hiroyuki Kato, Takahisa Ogasawara, Sho Ninomiya and Kaori Suzuki for technical assistance. We also thank Drs. Hajime Komano, Masato Yugami, and Tsuyoshi Matsuo for useful discussions.

---

**Supplemental Figure S1.**
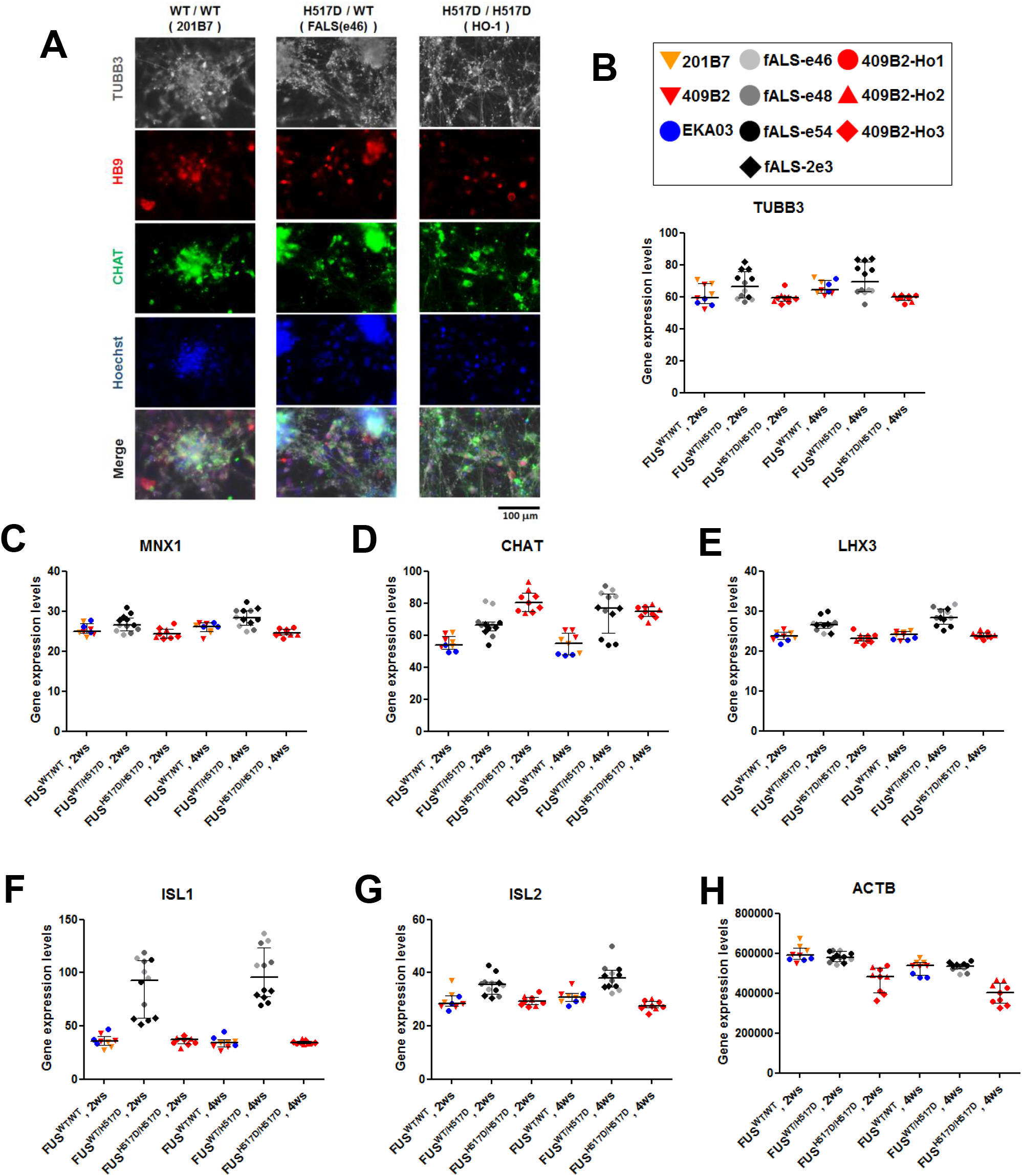
iPSC-derived MN differentiation. (A) Representative images of immunofluorescence of pan-neuron (TUBB3) and motor neuron (HB9 and CHAT) markers in MN terminal differentiated cells-derived from iPS cells. iPSC-derived MNs possessing FUSWT/WT, FUSWT/H517D or FUSH517D/H517D were fixed at 4 weeks after plating onto 96-well plates. (B-G) Microarray analyses of neuronal (TUBB3) and motor neuronal (MNX1/HB9, CHAT, LHX3 and ISL1/2) markers in expression. Values represent mean ± SD. (H) b-actin house keeping gene expression levels.

**Supplemental Figure S2.**
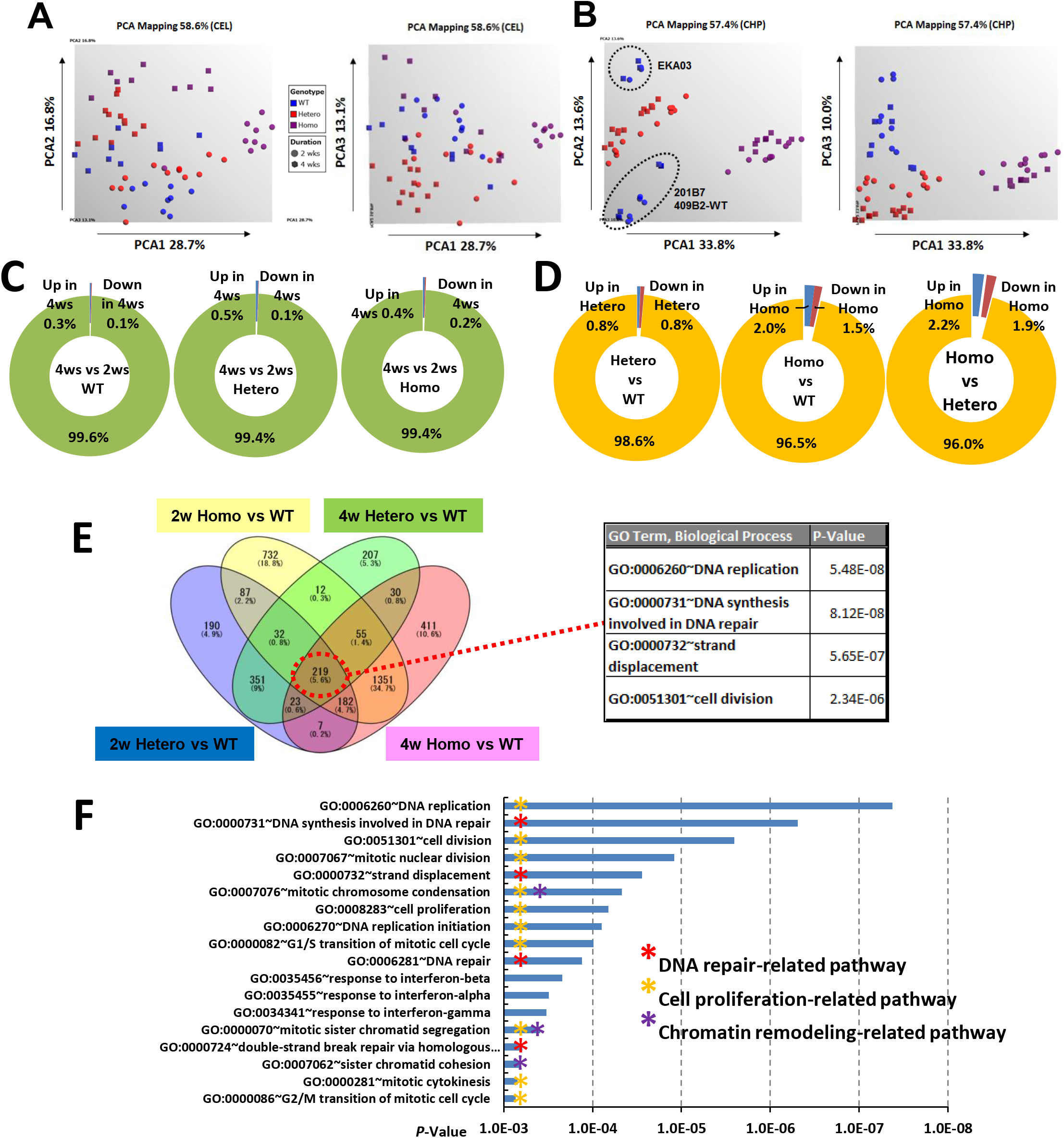
Large-scale transcriptome analysis with regular bioinformatics. (A, B) PCA analyses of large-scale transcriptome data. The graphics of the PCA are shown. Samples of iPSC-derived MNs possessing FUSWT/WT (WT), FUSWT/H517D (Hetero), and FUSH517D/H517D (Homo) are shown in blue, red, and purple, respectively. Samples differentiated from MN precursors for 2 weeks and 4 weeks were shown as spheres and squares, respectively. PCA analyses were performed with CEL files (A) and CHP files (B), respectively. (C) Populations of up-regulated (fold change (FC) >2 and down-regulated (FC < -2) DEGs in the 4ws MN differentiation relative to the 2ws differentiation. DEGs with FDR P-value < 0.05 were selected. (D) From the view point of the differences among FUS genotypes, population of DEGs between WT and Hetero groups, WT and Homo groups, and Hetero and Homo groups were shown. (E) Venn diagrams of DEGs were shown in the left figure. 219 common DEGs were applied for GO term analyses and top 4 GO terms were shown in the right table. (F) GO term analyses of down-regulated DEGs in Hetero groups relative to WT and top GO terms were shown (P-value < 0.001).

**Supplemental Figure S3.**
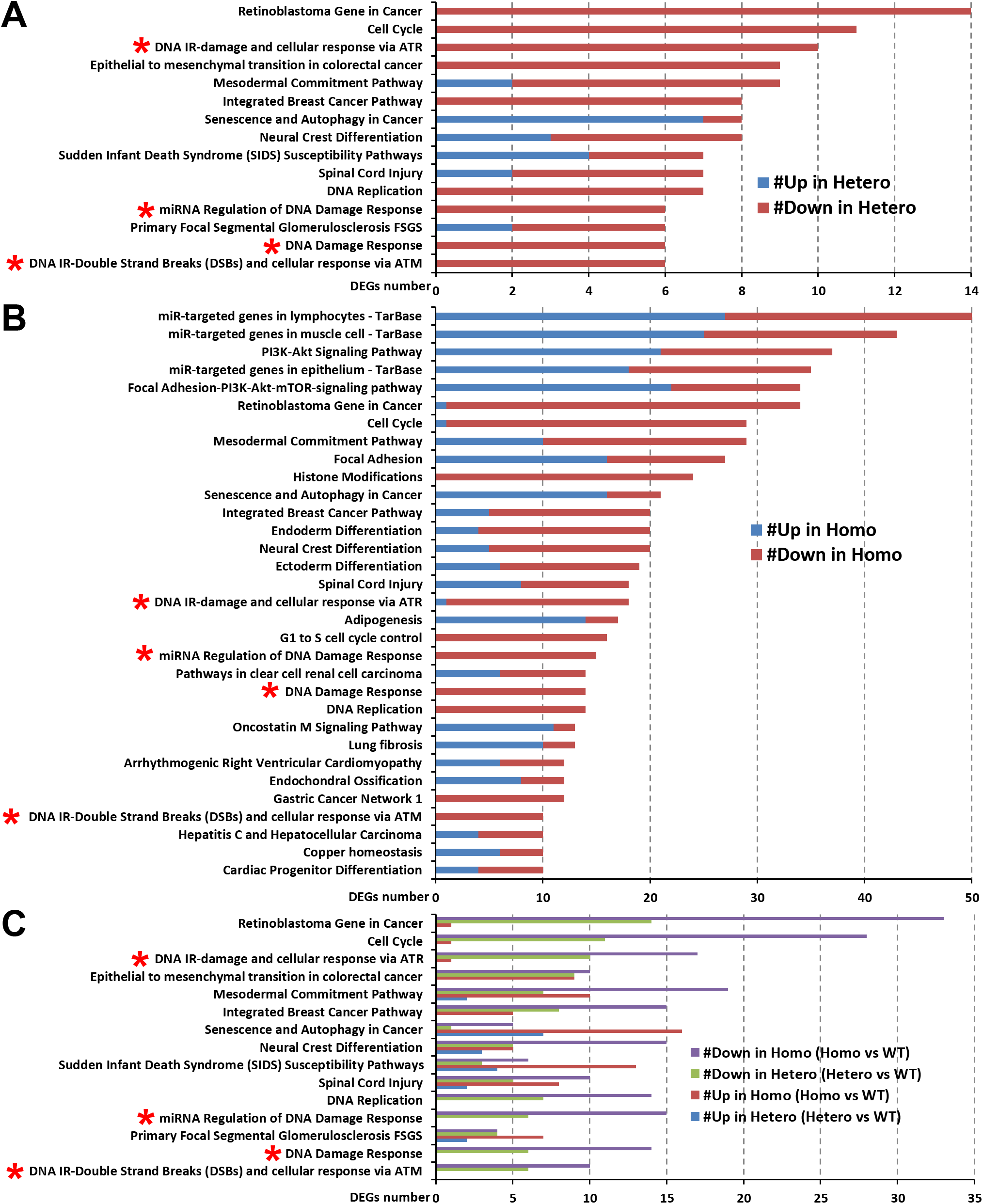
Pathway analysis with DEGs. (A)Pathway analysis was performed with DEGs between the Hetero and WT groups. Top pathways were shown in the graph (DEG number in each pathway _ 6, P-value < 0.05). The X-axis indicates the number of genes in each pathway which are shown on the Y-axis. Number of up-regulated DEGs were shown in blue and down-regulated DEGs were shown in red. The asterisks indicate “DNA repair”-related pathways. (B) Similarly, pathway analysis was performed with DEGs between the Homo and WT groups. Top pathways were shown in the graph (DEG number in each pathway _ 10, P-value < 0.001). (C) Top pathways found in the comparison between Hetero and WT were augmented in the comparison between Homo and WT. The asterisks indicate “DNA repair”-related pathways.

**Supplemental Figure S4.**
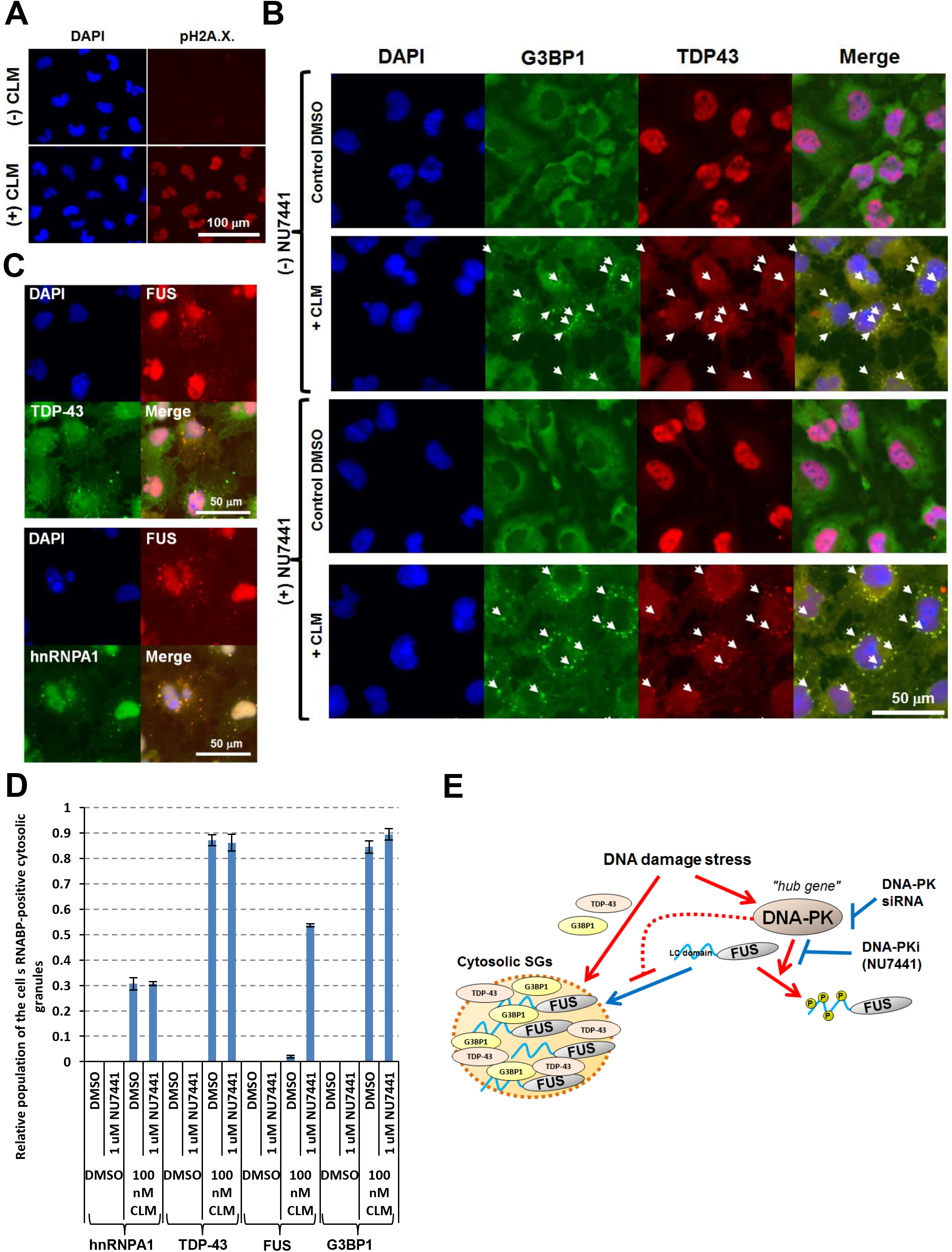
FUS is unique RNABP translocated to cytosolic SGs in a CLM- and DNA-PK inhibitor-dependent manner. (A) U251 MG (KO) cells were treated with CLM and the immunofluorescence imaging was performed with DAPI and anti-phospho-H2A.X. antibody for staining nuclei and DNA damages. (B, C) The cells were pre-treated with control DMSO or 1 mM NU7441, and then treated with control DMSO or CLM. Immunofluorescence imaging was performed. Arrowheads indicate the cytosolic granules where TDP-43 and G3BP1 were co-localized (B). Co-localization of FUS with TDP-43 and hnRNPA1 in the cytosolic granules was observed (C). (D) Four fields of the cells in each condition were imaged and the relative populations of cells possessing each RNABP-positive cytosolic SGs to the DAPI-positive cells were calculated. (E) A model for PRKDC in the regulation of cytosolic FUS mis-localization to SGs.

**Supplemental Figure S5.**
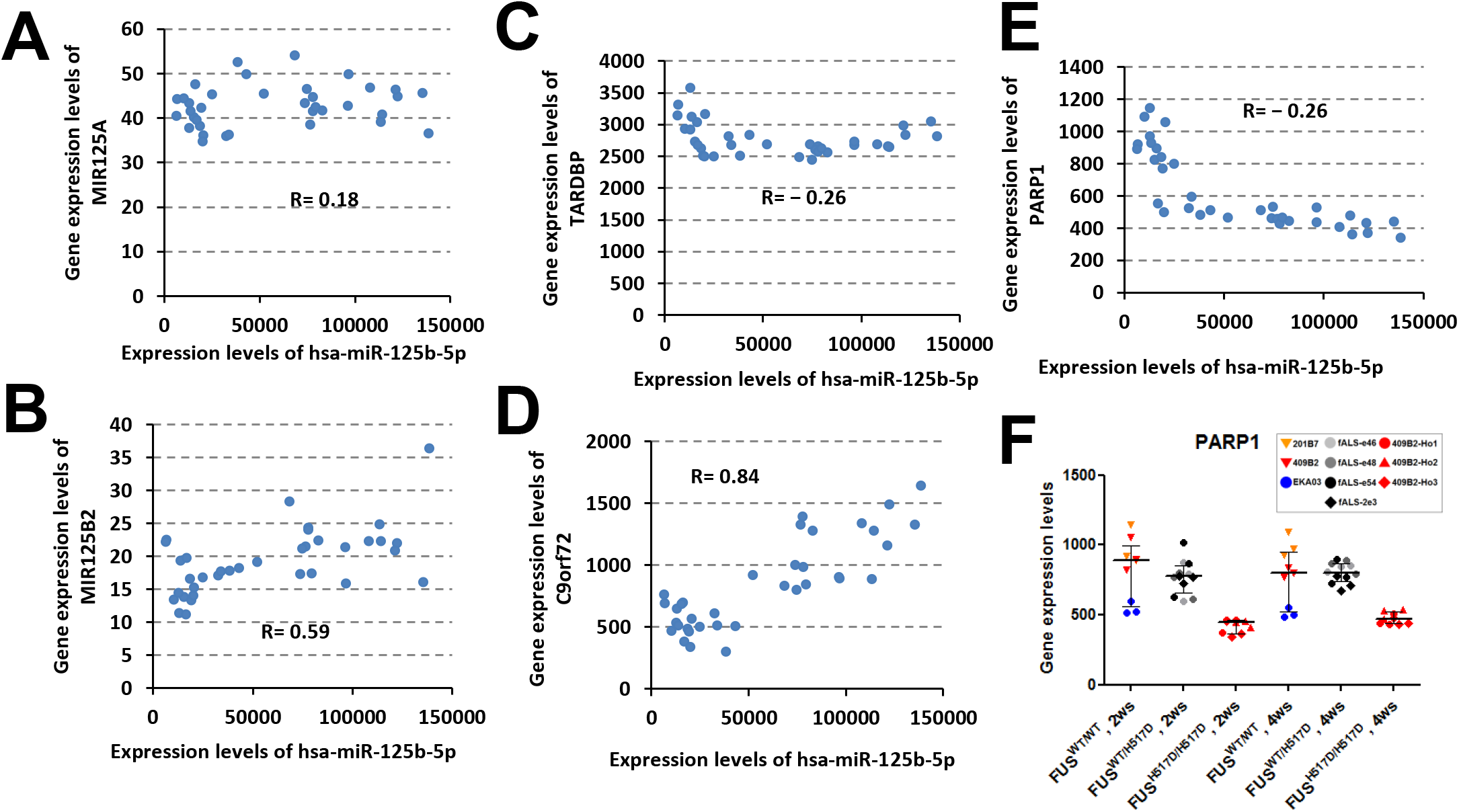
Correlation analyses of hsa-miR-125b-5p. (A, B) The expression levels of hsa-miR-125b-5p versus pri-miRNA transcripts in WT and Homo groups were plotted. (C-E) Correlation analyses of hsa-miR-125b-5p to TARDBP, C9orf72 and PARP1 mRNAs in WT and Homo groups. (F) PARP1 gene expression levels were down-regulated in the MN-Homo relative to the MN-WT. Values represent mean ± SD.

**Supplemental Figure S6.**
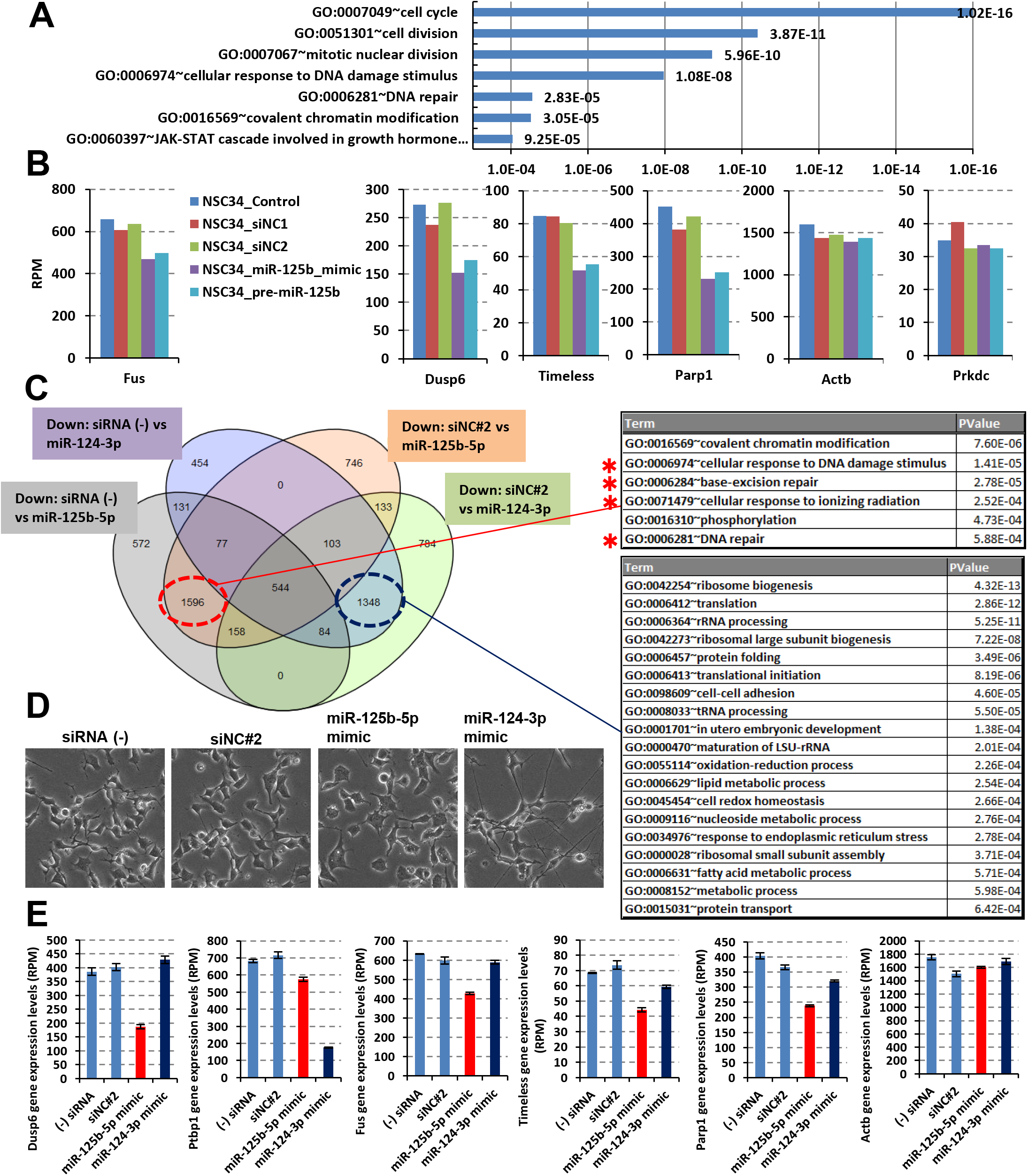
Introduction of miR-125b-5p into NSC-34 cells caused uniform down-regulation of the genes associating with “DNA repair”-related pathways. GO term analysis of down-regulated DEGs (FC < -1.5) in the NSC-34 cells transfected with miR-125b-5p mimic and pre-miR-125b in compared with siRNA (-), negative control siRNA #1 (siNC#1) and #2 (siNC#2) was performed. Top GO terms (Biological Process) were shown in the graph. (B) Gene expression levels provided by RNA-seq analyses. (C) Venn diagram of down-regulated DEGs (logFC < -0.5) in the cells treated with miR-125b-5p and miR-124-3p in comparison with siRNA (-) or siNC#2. Specific down-regulated 1596 DEGs by the introduction of miR-125b-5p were associated with “DNA repair”-related GO terms (shown in the upper table). On the other hand, specific down-regulated 1348 DEGs by the introduction of miR-124-3p were not associated with them (shown in the lower table). (D) Introduction of miR-124-3p mimic caused morphological changes with long processes. (E) Dusp6 and Ptbp1 mRNAs are known targets of miR-125b-5p and miR124-3p, respectively. Gene expression levels of Dusp6, Ptbp1, Fus, Timeless, Parp1 and Actb were shown. Values represent mean ± SD (n=3)

**Supplemental Table S1.**
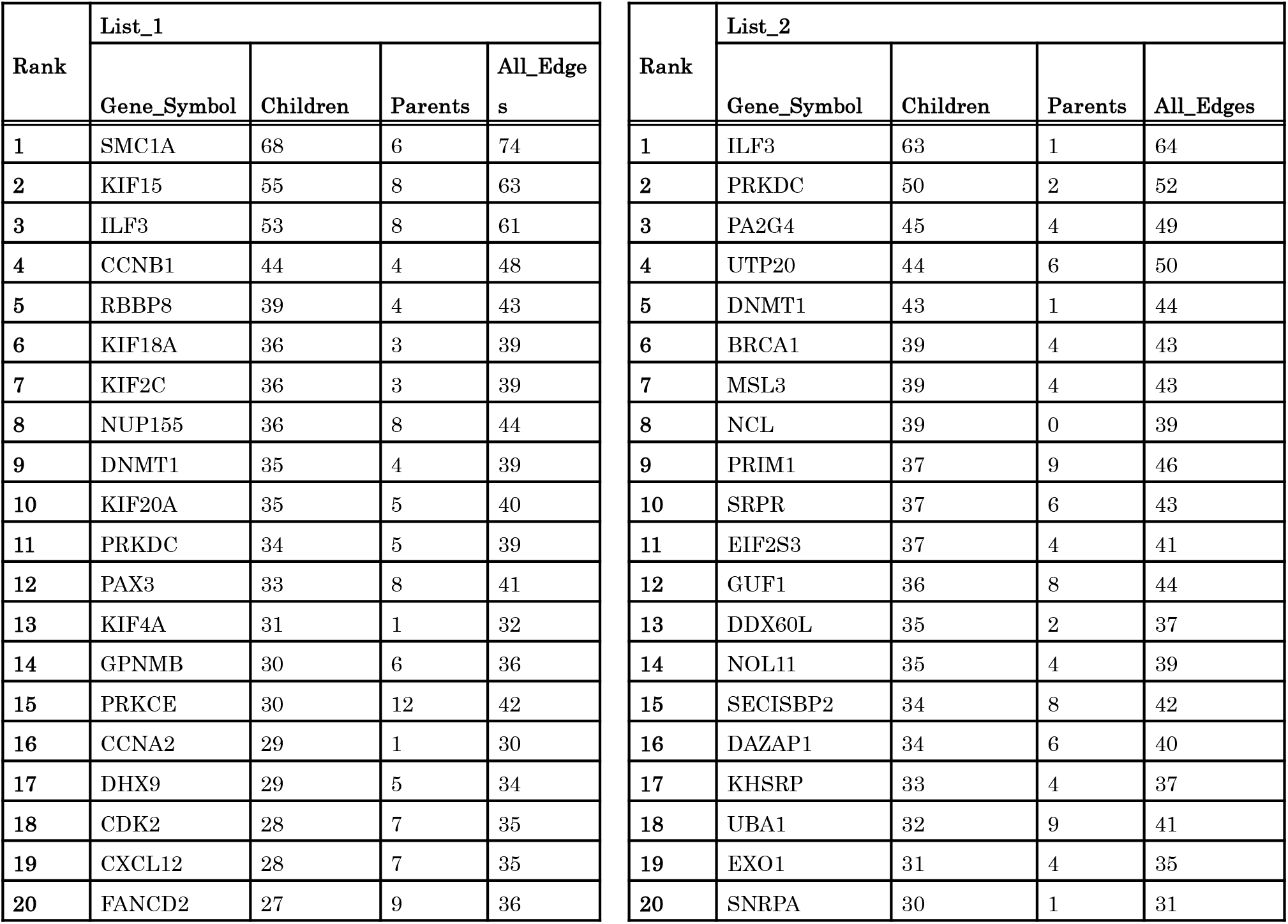
Top 20 “hub genes” in Bayesian gene regulatory network of ALS-related pathways and RBP genes. Top 20 “hub genes” in Bayesian gene regulatory network of ALS-related pathways and RBP genes are shown as List_1 and List_2, respectively. “Children” indicates the number of genes in the downstream of each gene in the calculated gene network. “Parents” indicates the number of genes in the upstream of each gene. “All_Edges” indicates the sum of the gene numbers in the downstream and upstream.

**Supplemental Table S2.** Top 20 “hub genes” in Bayesian gene regulatory network containing miRNAs. The top 20 “hub genes/miRNAs” in Bayesian GRN of ALS-related pathways and RBP genes with miRNAs are shown as List_3, and List_4, respectively.

| Rank | List_3 |  |  |  |
| --- | --- | --- | --- | --- |
|  | Gene_Symbol | Children | Parents | All_Edges |
| 1 | SMC1A | 52 | 4 | 56 |
| 2 | ILF3 | 40 | 3 | 43 |
| 3 | HNRNPM | 37 | 7 | 44 |
| 4 | KIF18A | 35 | 3 | 38 |
| 5 | hsa-miR-205-5p | 35 | 1 | 36 |
| 6 | hsa-miR-23b-3p | 31 | 4 | 35 |
| 7 | CCNB1 | 30 | 3 | 33 |
| 8 | SRSF3 | 30 | 11 | 41 |
| 9 | KIF15 | 29 | 5 | 34 |
| 10 | SNRPF | 29 | 11 | 40 |
| 11 | KIF20A | 28 | 4 | 32 |
| 12 | NUP205 | 27 | 4 | 31 |
| 13 | RAD23B | 27 | 6 | 33 |
| 14 | hsa-miR-125b-5p | 27 | 9 | 36 |
| 15 | RBBP8 | 26 | 4 | 30 |
| 16 | NCL | 25 | 3 | 28 |
| 17 | PPARGC1A | 25 | 6 | 31 |
| 18 | POLR2E | 24 | 6 | 30 |
| 19 | SESN3 | 23 | 10 | 33 |
| 20 | CCND2 | 23 | 8 | 31 |

| Rank | List_4 |  |  |  |
| --- | --- | --- | --- | --- |
|  | Gene_Symbol | Children | Parents | All_Edges |
| 1 | ILF3 | 43 | 2 | 45 |
| 2 | MSL3 | 42 | 5 | 47 |
| 3 | DAZAP1 | 41 | 4 | 45 |
| 4 | UTP20 | 38 | 8 | 46 |
| 5 | NCL | 37 | 4 | 41 |
| 6 | HNRNPM | 37 | 5 | 42 |
| 7 | IGF2BP1 | 36 | 7 | 43 |
| 8 | RPLP0 | 35 | 2 | 37 |
| 9 | NOL11 | 35 | 1 | 36 |
| 10 | EIF2AK3 | 31 | 4 | 35 |
| 11 | PRKDC | 30 | 5 | 35 |
| 12 | SNRPF | 30 | 9 | 39 |
| 13 | EWSR1 | 29 | 3 | 32 |
| 14 | hsa-miR-205-5p | 29 | 1 | 30 |
| 15 | GUF1 | 28 | 5 | 33 |
| 16 | hsa-miR-125b-5p | 28 | 12 | 40 |
| 17 | POLR2E | 27 | 5 | 32 |
| 18 | EXOSC10 | 26 | 10 | 36 |
| 19 | EIF2S3 | 26 | 6 | 32 |
| 20 | PRIM1 | 26 | 1 | 27 |

**Supplemental Table S3.**
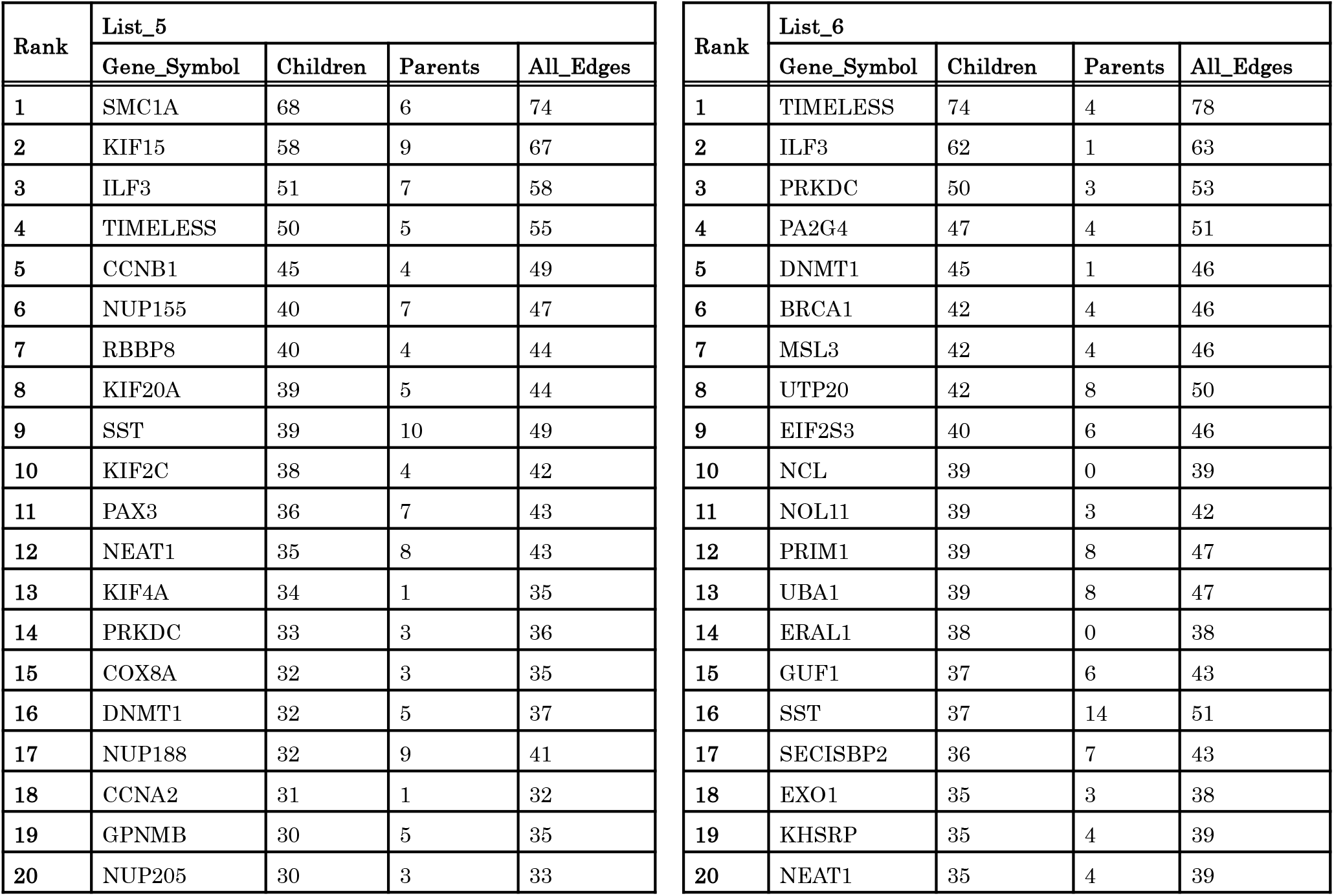
Top 20 “hub genes” in Bayesian gene regulatory network with additional AD-related and circadian rhythm-related genes. Top 20 “hub genes” in the regulatory gene networks of ALS-related pathways and RBP genes with additional AD-related and circadian rhythm-related genes are shown as List 5 and List 6, respectively.

**Supplemental Table S4.**
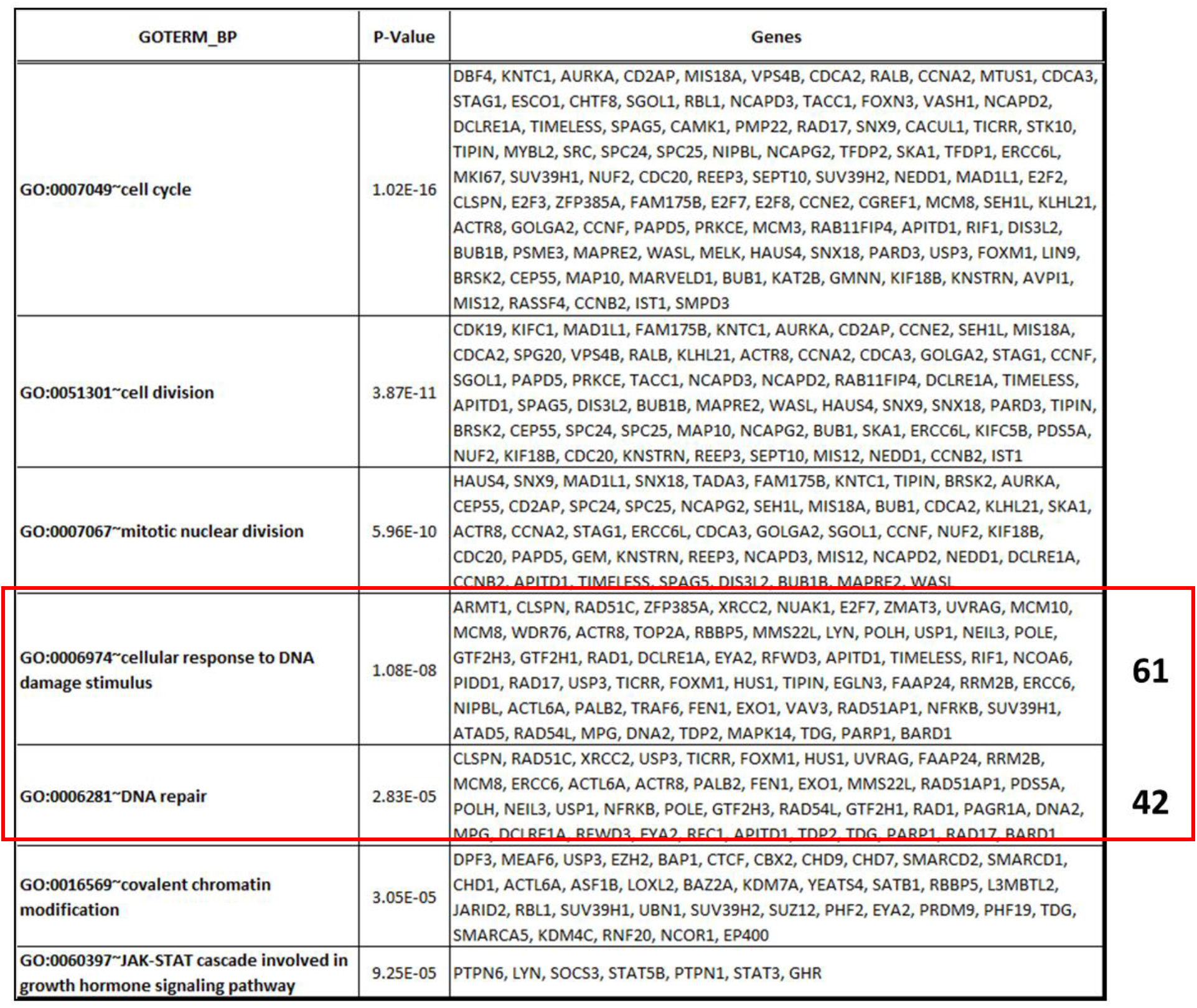
Top pathways associating with down-regulated DEGs in the NSC-34 cells transiently transfected with miR-125b mimic and pre-miR-125b. Down-regulated DEGs (logFC < -0.5, P-value < 0.05) in the NSC-34 cell transiently transfected with miR-125b mimic and pre-miR-125b were selected for GO term analyses. “cellular response to DNA damage stimulus” and “DNA repair” have 61 and 42 memberships.

**Supplemental Table S5.**
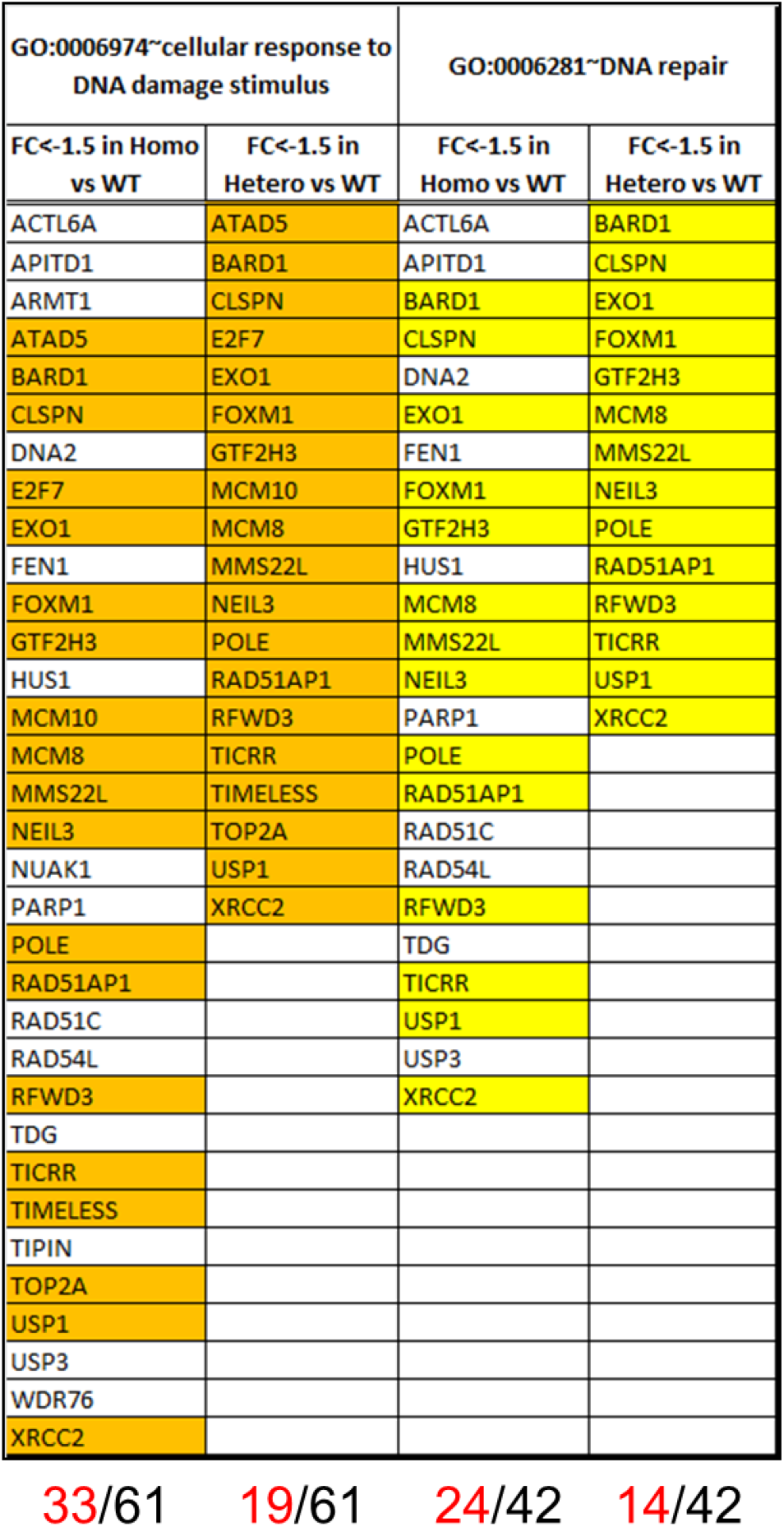
“DNA repair”-related down-regulated DEGs in the NSC-34 cells transiently transfected with miR-125b mimic and pre-miR-125b showed lower expression levels in the FUSH517D MNs than in the FUSWT MNs. In the 61 down-regulated DEGs of the GO term “cellular response to DNA damage stimulus”, 19 down-regulated DEGs showed lower expression levels (FC < -1.5) in Hetero than WT groups. In addition, 33 down-regulated DEGs showed lower expression levels (FC < -1.5) in Homo than WT groups. Shared down-regulated DEGs between Hetero vs WT and Homo vs WT are shown in orange. On the other hand, in the 42 down-regulated DEGs of the GO term “DNA repair”, 14 down-regulated DEGs showed lower expression levels (FC < -1.5) in Hetero group than in the WT groups. Furthermore, 24 down-regulated DEGs showed lower expression levels (FC <-1.5) in Homo group than in the WT groups. Similarly, shared down-regulated DEGs between Hetero vs WT and Homo vs WT were shown in yellow.

| GO:0006974~cellular response to DNA damage stimulus |  | GO:0006281~DNA repair |  |
| --- | --- | --- | --- |
| FC<-1.5 in Homo vs WT | FC<-1.5 in Hetero vs WT | FC<-1.5 in Homo vs WT | FC<-1.5 in Hetero vs WT |
| ACTL6A | ATAD5 | ACTL6A | BARD1 |
| APITD1 | BARD1 | APITD1 | CLSPN |
| ARMT1 | CLSPN | BARD1 | EXO1 |
| ATAD5 | E2F7 | CLSPN | FOXM1 |
| BARD1 | EXO1 | DNA2 | GTF2H3 |
| CLSPN | FOXM1 | EXO1 | MCM8 |
| DNA2 | GTF2H3 | FEN1 | MMS22L |
| E2F7 | MCM10 | FOXM1 | NEIL3 |
| EXO1 | MCM8 | GTF2H3 | POLE |
| FEN1 | MMS22L | HUS1 | RAD51AP1 |
| FOXM1 | NEIL3 | MCM8 | RFWD3 |
| GTF2H3 | POLE | MMS22L | TICRR |
| HUS1 | RAD51AP1 | NEIL3 | USP1 |
| MCM10 | RFWD3 | PARP1 | XRCC2 |
| MCM8 | TICRR | POLE |  |
| MMS22L | TIMELESS | RAD51AP1 |  |
| NEIL3 | TOP2A | RAD51C |  |
| NUAK1 | USP1 | RAD54L |  |
| PARP1 | XRCC2 | RFWD3 |  |
| POLE |  | TDG |  |
| RAD51AP1 |  | TICRR |  |
| RAD51C |  | USP1 |  |
| RAD54L |  | USP3 |  |
| RFWD3 |  | XRCC2 |  |
| TDG |  |  |  |
| TICRR |  |  |  |
| TIMELESS |  |  |  |
| TIPIN |  |  |  |
| TOP2A |  |  |  |
| USP1 |  |  |  |
| USP3 |  |  |  |
| WDR76 |  |  |  |
| XRCC2 |  |  |  |

## REFFERENCES

Affara, M., Dunmore, B., Savoie, C., Imoto, S., Tamada, Y., Araki, H., Charnock-Jones, D.S., Miyano, S., and Print, C. (2007) Understanding endothelial cell apoptosis: what can the transcriptome, glycome and proteome reveal. Philos Trans R Soc Lond B Biol Sci, 362, 1469–1487.

Akiyama, T., Suzuki, N., Ishikawa, M., Fujimori, K., Sone, T., Kawada, J., Funayama, R., Fujishima, F., Mitsuzawa, S., Ikeda, K. et al. (2019) Aberrant axon branching via Fos-B dysregulation in FUS-ALS motor neurons. EBioMedicine, 45, 362–378.

Anderson, P., and Kedersha, N. (2009) RNA granules: post-transcriptional and epigenetic modulators of gene expression. Nat Rev Mol Cell Biol, 10, 430–436.

Aulas, A., Stabile, S., and Vande Velde, C. (2012) Endogenous TDP-43, but not FUS, contributes to stress granule assembly via G3BP. Mol Neurodegener, 7, 54.

Balakrishnan, I., Yang, X., Brown, J., Ramakrishnan, A., Torok-Storb, B., Kabos, P., Hesselberth, J.R., and Pillai, M.M. (2014) Genome-wide analysis of miRNA-mRNA interactions in marrow stromal cells. Stem Cells, 32, 662–673.

Banzhaf-Strathmann, J., Benito, E., May, S., Arzberger, T., Tahirovic, S., Kretzschmar, H., Fischer, A., and Edbauer, D. (2014) MicroRNA-125b induces tau hyperphosphorylation and cognitive deficits in Alzheimer’s disease. EMBO J, 33, 1667–1680.

Barmada, S.J. (2015) Linking RNA Dysfunction and Neurodegeneration in Amyotrophic Lateral Sclerosis. Neurotherapeutics, 12, 340–351.

Bentmann, E., Neumann, M., Tahirovic, S., Rodde, R., Dormann, D., and Haass, C. (2012) Requirements for stress granule recruitment of fused in sarcoma (FUS) and TAR DNA-binding protein of 43 kDa (TDP-43). J Biol Chem, 287, 23079–23094.

Bosco, D.A., Lemay, N., Ko, H.K., Zhou, H., Burke, C., Kwiatkowski, T.J., Sapp, P., McKenna-Yasek, D., Brown, R.H., and Hayward, L.J. (2010) Mutant FUS proteins that cause amyotrophic lateral sclerosis incorporate into stress granules. Hum Mol Genet, 19, 4160–4175.

Brown, R.H., and Al-Chalabi, A. (2017) Amyotrophic Lateral Sclerosis. N Engl J Med, 377, 162–172.

Cardinale, A., Racaniello, M., Saladini, S., De Chiara, G., Mollinari, C., de Stefano, M.C., Pocchiari, M., Garaci, E., and Merlo, D. (2012) Sublethal doses of β-amyloid peptide abrogate DNA-dependent protein kinase activity. J Biol Chem, 287, 2618–2631.

Chaudhuri, A.A., So, A.Y., Mehta, A., Minisandram, A., Sinha, N., Jonsson, V.D., Rao, D.S., O’Connell, R.M., and Baltimore, D. (2012) Oncomir miR-125b regulates hematopoiesis by targeting the gene Lin28A. Proc Natl Acad Sci U S A, 109, 4233–4238.

Dash, S., Balasubramaniam, M., Rana, T., Godino, A., Peck, E.G., Goodwin, J.S., Villalta, F., Calipari, E.S., Nestler, E.J., Dash, C. et al. (2017) Poly (ADP-Ribose) Polymerase-1 (PARP-1) Induction by Cocaine Is Post-Transcriptionally Regulated by miR-125b. eNeuro, 4,

Davydov, V., Hansen, L.A., and Shackelford, D.A. (2003) Is DNA repair compromised in Alzheimer’s disease. Neurobiol Aging, 24, 953–968.

DeJesus-Hernandez, M., Kocerha, J., Finch, N., Crook, R., Baker, M., Desaro, P., Johnston, A., Rutherford, N., Wojtas, A., Kennelly, K. et al. (2010) De novo truncating FUS gene mutation as a cause of sporadic amyotrophic lateral sclerosis. Hum Mutat, 31, E1377–89.

Deng, Q., Holler, C.J., Taylor, G., Hudson, K.F., Watkins, W., Gearing, M., Ito, D., Murray, M.E., Dickson, D.W., Seyfried, N.T. et al. (2014) FUS is phosphorylated by DNA-PK and accumulates in the cytoplasm after DNA damage. J Neurosci, 34, 7802–7813.

Dormann, D., and Haass, C. (2011) TDP-43 and FUS: a nuclear affair. Trends Neurosci, 34, 339–348.

Dormann, D., Rodde, R., Edbauer, D., Bentmann, E., Fischer, I., Hruscha, A., Than, M.E., Mackenzie, I.R., Capell, A., Schmid, B. et al. (2010) ALS-associated fused in sarcoma (FUS) mutations disrupt Transportin-mediated nuclear import. EMBO J, 29, 2841–2857.

Rj, E.B.A.T. (1994) An Introduction to the Bootstrap. Boca Raton, FL, CRC Press.,

Ferrante, R.J., Browne, S.E., Shinobu, L.A., Bowling, A.C., Baik, M.J., MacGarvey, U., Kowall, N.W., Brown, R.H., and Beal, M.F. (1997) Evidence of increased oxidative damage in both sporadic and familial amyotrophic lateral sclerosis. J Neurochem, 69, 2064–2074.

Friedman, N., Linial, M., Nachman, I., and Pe’er, D. (2000) Using Bayesian networks to analyze expression data. J Comput Biol, 7, 601–620.

Fujimori, K., Ishikawa, M., Otomo, A., Atsuta, N., Nakamura, R., Akiyama, T., Hadano, S., Aoki, M., Saya, H., Sobue, G. et al. (2018) Modeling sporadic ALS in iPSC-derived motor neurons identifies a potential therapeutic agent. Nat Med,

Gama-Carvalho, M., L Garcia-Vaquero, M., R Pinto, F., Besse, F., Weis, J., Voigt, A., Schulz, J.B., and De Las Rivas, J. (2017) Linking amyotrophic lateral sclerosis and spinal muscular atrophy through RNA-transcriptome homeostasis: a genomics perspective. J Neurochem, 141, 12–30.

Gao, P., Yoo, S.H., Lee, K.J., Rosensweig, C., Takahashi, J.S., Chen, B.P., and Green, C.B. (2013) Phosphorylation of the cryptochrome 1 C-terminal tail regulates circadian period length. J Biol Chem, 288, 35277–35286.

Gerstberger, S., Hafner, M., and Tuschl, T. (2014) A census of human RNA-binding proteins. Nat Rev Genet, 15, 829–845.

Hafner, M., Landthaler, M., Burger, L., Khorshid, M., Hausser, J., Berninger, P., Rothballer, A., Ascano, M.J., Jungkamp, A.C., Munschauer, M. et al. (2010) Transcriptome-wide identification of RNA-binding protein and microRNA target sites by PAR-CLIP. Cell, 141, 129–141.

Han, T.W., Kato, M., Xie, S., Wu, L.C., Mirzaei, H., Pei, J., Chen, M., Xie, Y., Allen, J., Xiao, G. et al. (2012) Cell-free formation of RNA granules: bound RNAs identify features and components of cellular assemblies. Cell, 149, 768–779.

Hartemink, A.J., Gifford, D.K., Jaakkola, T.S., and Young, R.A. (2001) Using graphical models and genomic expression data to statistically validate models of genetic regulatory networks. Pac Symp Biocomput, 422–433.

Huynh-Thu, V.A., Irrthum, A., Wehenkel, L., and Geurts, P. (2010) Inferring regulatory networks from expression data using tree-based methods. PLoS One, 5,

Ichiyanagi, N., Fujimori, K., Yano, M., Ishihara-Fujisaki, C., Sone, T., Akiyama, T., Okada, Y., Akamatsu, W., Matsumoto, T., Ishikawa, M. et al. (2016) Establishment of In Vitro FUS-Associated Familial Amyotrophic Lateral Sclerosis Model Using Human Induced Pluripotent Stem Cells. Stem Cell Reports,

Imaizumi, K., Sone, T., Ibata, K., Fujimori, K., Yuzaki, M., Akamatsu, W., and Okano, H. (2015) Controlling the Regional Identity of hPSC-Derived Neurons to Uncover Neuronal Subtype Specificity of Neurological Disease Phenotypes. Stem Cell Reports, 5, 1010–1022.

Imoto, S., Goto, T., and Miyano, S. (2002) Estimation of genetic networks and functional structures between genes by using Bayesian networks and nonparametric regression. Pac Symp Biocomput, 175–186.

Izumi, R., Warita, H., Niihori, T., Takahashi, T., Tateyama, M., Suzuki, N., Nishiyama, A., Shirota, M., Funayama, R., Nakayama, K. et al. (2015) Isolated inclusion body myopathy caused by a multisystem proteinopathy-linked hnRNPA1 mutation. Neurol Genet, 1, e23.

Kanungo, J. (2013) DNA-dependent protein kinase and DNA repair: relevance to Alzheimer’s disease. Alzheimers Res Ther, 5, 13.

Karagkouni, D., Paraskevopoulou, M.D., Chatzopoulos, S., Vlachos, I.S., Tastsoglou, S., Kanellos, I., Papadimitriou, D., Kavakiotis, I., Maniou, S., Skoufos, G. et al. (2018) DIANA-TarBase v8: a decade-long collection of experimentally supported miRNA-gene interactions. Nucleic Acids Res, 46, D239–D245.

Kim, H.J., Kim, N.C., Wang, Y.D., Scarborough, E.A., Moore, J., Diaz, Z., MacLea, K.S., Freibaum, B., Li, S., Molliex, A. et al. (2013) Mutations in prion-like domains in hnRNPA2B1 and hnRNPA1 cause multisystem proteinopathy and ALS. Nature, 495, 467–473.

Lattante, S., Rouleau, G.A., and Kabashi, E. (2013) TARDBP and FUS mutations associated with amyotrophic lateral sclerosis: summary and update. Hum Mutat, 34, 812–826.

Lenzi, J., De Santis, R., de Turris, V., Morlando, M., Laneve, P., Calvo, A., Caliendo, V., Chiò, A., Rosa, A., and Bozzoni, I. (2015) ALS mutant FUS proteins are recruited into stress granules in induced pluripotent stem cell-derived motoneurons. Dis Model Mech, 8, 755–766.

Liu, X., Chen, J., Liu, W., Li, X., Chen, Q., Liu, T., Gao, S., and Deng, M. (2015) The fused in sarcoma protein forms cytoplasmic aggregates in motor neurons derived from integration-free induced pluripotent stem cells generated from a patient with familial amyotrophic lateral sclerosis carrying the FUS-P525L mutation. Neurogenetics, 16, 223–231.

Ludwig, N., Leidinger, P., Becker, K., Backes, C., Fehlmann, T., Pallasch, C., Rheinheimer, S., Meder, B., Stähler, C., Meese, E. et al. (2016) Distribution of miRNA expression across human tissues. Nucleic Acids Res, 44, 3865–3877.

Ma, X., Liu, L., and Meng, J. (2017) MicroRNA-125b promotes neurons cell apoptosis and Tau phosphorylation in Alzheimer’s disease. Neurosci Lett, 661, 57–62.

Margolin, A.A., Nemenman, I., Basso, K., Wiggins, C., Stolovitzky, G., Dalla Favera, R., and Califano, A. (2006) ARACNE: an algorithm for the reconstruction of gene regulatory networks in a mammalian cellular context. BMC Bioinformatics, 7 Suppl 1, S7.

Mazzoccoli, G., Laukkanen, M.O., Vinciguerra, M., Colangelo, T., and Colantuoni, V. (2016) A Timeless Link Between Circadian Patterns and Disease. Trends Mol Med, 22, 68–81.

Monahan, Z., Ryan, V.H., Janke, A.M., Burke, K.A., Rhoads, S.N., Zerze, G.H., O’Meally, R., Dignon, G.L., Conicella, A.E., Zheng, W. et al. (2017) Phosphorylation of the FUS low-complexity domain disrupts phase separation, aggregation, and toxicity. EMBO J,

Morlando, M., Dini Modigliani, S., Torrelli, G., Rosa, A., Di Carlo, V., Caffarelli, E., and Bozzoni, I. (2012) FUS stimulates microRNA biogenesis by facilitating co-transcriptional Drosha recruitment. EMBO J, 31, 4502–4510.

Murray, D.T., Kato, M., Lin, Y., Thurber, K.R., Hung, I., McKnight, S.L., and Tycko, R. (2017) Structure of FUS Protein Fibrils and Its Relevance to Self-Assembly and Phase Separation of Low-Complexity Domains. Cell,

Musiek, E.S., and Holtzman, D.M. (2016) Mechanisms linking circadian clocks, sleep, and neurodegeneration. Science, 354, 1004–1008.

Naujock, M., Stanslowsky, N., Bufler, S., Naumann, M., Reinhardt, P., Sterneckert, J., Kefalakes, E., Kassebaum, C., Bursch, F., Lojewski, X. et al. (2016) 4-Aminopyridine Induced Activity Rescues Hypoexcitable Motor Neurons from Amyotrophic Lateral Sclerosis Patient-Derived Induced Pluripotent Stem Cells. Stem Cells, 34, 1563–1575.

Naumann, M., Pal, A., Goswami, A., Lojewski, X., Japtok, J., Vehlow, A., Naujock, M., Günther, R., Jin, M., Stanslowsky, N. et al. (2018) Impaired DNA damage response signaling by FUS-NLS mutations leads to neurodegeneration and FUS aggregate formation. Nat Commun, 9, 335.

Neumann, M., Rademakers, R., Roeber, S., Baker, M., Kretzschmar, H.A., and Mackenzie, I.R. (2009) A new subtype of frontotemporal lobar degeneration with FUS pathology. Brain, 132, 2922–2931.

Parisi, C., Napoli, G., Amadio, S., Spalloni, A., Apolloni, S., Longone, P., and Volonté, C. (2016) MicroRNA-125b regulates microglia activation and motor neuron death in ALS. Cell Death Differ, 23, 531–541.

Pillai, M.M., Gillen, A.E., Yamamoto, T.M., Kline, E., Brown, J., Flory, K., Hesselberth, J.R., and Kabos, P. (2014) HITS-CLIP reveals key regulators of nuclear receptor signaling in breast cancer. Breast Cancer Res Treat, 146, 85–97.

Rhoads, S.N., Monahan, Z.T., Yee, D.S., Leung, A.Y., Newcombe, C.G., O’Meally, R.N., Cole, R.N., and Shewmaker, F.P. (2018) The prionlike domain of FUS is multiphosphorylated following DNA damage without altering nuclear localization. Mol Biol Cell, 29, 1786–1797.

Suzuki, N., Kato, S., Kato, M., Warita, H., Mizuno, H., Kato, M., Shimakura, N., Akiyama, H., Kobayashi, Z., Konno, H. et al. (2012) FUS/TLS-immunoreactive neuronal and glial cell inclusions increase with disease duration in familial amyotrophic lateral sclerosis with an R521C FUS/TLS mutation. J Neuropathol Exp Neurol, 71, 779–788.

Tamada, Y., Shimamura, T., Yamaguchi, R., Imoto, S., Nagasaki, M., and Miyano, S. (2011) Sign: large-scale gene network estimation environment for high performance computing. Genome Inform, 25, 40–52.

Taylor, J.P., Brown, R.H., and Cleveland, D.W. (2016) Decoding ALS: from genes to mechanism. Nature, 539, 197–206.

Tyzack, G.E., Luisier, R., Taha, D.M., Neeves, J., Modic, M., Mitchell, J.S., Meyer, I., Greensmith, L., Newcombe, J., Ule, J. et al. (2019) Widespread FUS mislocalization is a molecular hallmark of amyotrophic lateral sclerosis. Brain, 142, 2572–2580.

Vance, C., Rogelj, B., Hortobagyi, T., De Vos, K.J., Nishimura, A.L., Sreedharan, J., Hu, X., Smith, B., Ruddy, D., Wright, P. et al. (2009) Mutations in FUS, an RNA processing protein, cause familial amyotrophic lateral sclerosis type 6. Science, 323, 1208–1211.

Vance, C., Scotter, E.L., Nishimura, A.L., Troakes, C., Mitchell, J.C., Kathe, C., Urwin, H., Manser, C., Miller, C.C., Hortobágyi, T. et al. (2013) ALS mutant FUS disrupts nuclear localization and sequesters wild-type FUS within cytoplasmic stress granules. Hum Mol Genet, 22, 2676–2688.

H, V. (2005) Bootstrap Tutorial. Mathematica Journal, 9, 768–775.

Wang, H., Guo, W., Mitra, J., Hegde, P.M., Vandoorne, T., Eckelmann, B.J., Mitra, S., Tomkinson, A.E., Van Den Bosch, L., and Hegde, M.L. (2018) Mutant FUS causes DNA ligation defects to inhibit oxidative damage repair in Amyotrophic Lateral Sclerosis. Nat Commun, 9, 3683.

Wang, W.Y., Pan, L., Su, S.C., Quinn, E.J., Sasaki, M., Jimenez, J.C., Mackenzie, I.R., Huang, E.J., and Tsai, L.H. (2013) Interaction of FUS and HDAC1 regulates DNA damage response and repair in neurons. Nat Neurosci, 16, 1383–1391.

Xie, S., Mortusewicz, O., Ma, H.T., Herr, P., Poon, R.Y., Poon, R.R., Helleday, T., and Qian, C. (2015) Timeless Interacts with PARP-1 to Promote Homologous Recombination Repair. Mol Cell, 60, 163–176.

Young, L.M., Marzio, A., Perez-Duran, P., Reid, D.A., Meredith, D.N., Roberti, D., Star, A., Rothenberg, E., Ueberheide, B., and Pagano, M. (2015) TIMELESS Forms a Complex with PARP1 Distinct from Its Complex with TIPIN and Plays a Role in the DNA Damage Response. Cell Rep, 13, 451–459.

Zhang, B., Gaiteri, C., Bodea, L.G., Wang, Z., McElwee, J., Podtelezhnikov, A.A., Zhang, C., Xie, T., Tran, L., Dobrin, R. et al. (2013) Integrated systems approach identifies genetic nodes and networks in late-onset Alzheimer’s disease. Cell, 153, 707–720.

